# Allosteric Inhibition of NDM-1 by Thanatin Preserves the Di-Zinc Center While Restricting Dynamics

**DOI:** 10.64898/2026.02.25.707299

**Authors:** Gwladys Rivière, Prince Kumar, Thomas Cummins, Ansel Hsiao, Leonard J. Mueller

**Affiliations:** Department of Chemistry, University of California – Riverside, Riverside, CA 92521, United States of America; Department of Radiation Oncology, David Geffen School of Medicine, University of California –Los Angeles, Los Angeles, CA 90024, United States of America; Department of Microbiology & Plant Pathology, University of California Riverside, Riverside, CA 92521, USA

## Abstract

The New Delhi metallo-β-lactamase 1 (NDM-1) is a major driver of carbapenem resistance in Gram-negative pathogens, yet the molecular basis by which antimicrobial peptides inhibit this enzyme has remained unresolved. Thanatin, a disulfide-stabilized β-hairpin peptide, was previously proposed to inactivate NDM-1 by displacing catalytic Zn²⁺ ions, but this model lacked direct structural support. Here, we combine high-resolution NMR spectroscopy, intermolecular NOE mapping, HADDOCK-guided docking, and molecular dynamics simulations to reveal a distinct zinc-retaining dynamic allosteric mechanism. Thanatin binds adjacent to the catalytic groove, preserving the native di-zinc coordination environment while simultaneously rigidifying the L3 catalytic loop, as confirmed by Zn-bound spectral fingerprints and EDTA titration experiments. This conformational restriction explains how the peptide inhibits the enzyme while maintaining a zinc-bound but catalytically compromised state, a finding that contrasts with zinc-displacement hypotheses and reconciles prior biochemical observations with structural data. In bacterial assays, this allosteric inhibition translates to a moderate restoration of carbapenem sensitivity, resulting in a 50% reduction in viable cell output even under high-level enzyme expression. Our findings establish a mechanistic framework for designing next-generation peptide inhibitors that target the dynamic vulnerabilities of metallo-β-lactamases.

## INTRODUCTION

Antimicrobial resistance (AMR) poses an escalating global threat^1^, with multidrug-resistant (MDR) Gram-negative pathogens driving rising morbidity and mortality^2^. Among the most clinically important resistance factors is New Delhi metallo-β-lactamase 1 (NDM-1), a broad-spectrum carbapenemase that hydrolyzes nearly all β-lactam antibiotics^3,4^. Anchored to the bacterial outer membrane through an N-terminal lipidation signal, NDM-1 contributes both enzymatic resistance and rapid horizontal dissemination of carbapenemase genes^5,6^. Reflecting its clinical importance, the World Health Organization has designated NDM-1-producing Enterobacteriaceae as top-priority targets for new antimicrobial development^7,8^.

Thanatin, a 21-residue disulfide-stabilized β-hairpin peptide from *Podisus maculiventris* exhibits potent activity against NDM-1–producing strains^5^. Its amphipathic fold enables engagement of both membrane-associated Lpt components (LptA, LptC, LptD)^6,9^ and the metallo-β-lactamase NDM-1^5^. Thanatin binds Lpt proteins with nanomolar affinity and NDM-1 with micromolar affinity, restoring carbapenem susceptibility in resistant isolates^5,6^. Despite these properties, the molecular mechanism by which thanatin inhibits NDM-1 remains unclear.

A zinc-displacement model has been previously proposed, suggesting that thanatin releases the catalytic Zn²⁺ ions of NDM-1^5^, but this hypothesis has lacked direct structural validation. Later studies highlighted the importance of the β-hairpin fold without clarifying how the peptide engages the enzyme^10^. The L3 loop modulates substrate access to the catalytic groove through its flexible motions^4,11^. Disrupting these dynamics could provide an alternative inhibition strategy that does not require metal removal. The absence of structural and dynamic information has limited rational design of peptide-based inhibitors targeting metallo-β-lactamases.

Here, we combine high-resolution NMR spectroscopy, HADDOCK-guided docking, and molecular dynamics simulations to define the structural and dynamic basis of NDM-1 inhibition by thanatin. Our data show that thanatin binds adjacent to the catalytic groove while preserving the native di-zinc coordination environment. Rather than releasing metal ions, the peptide engages residues surrounding the flexible L3 loop and selectively suppresses its intrinsic motions. This dynamics-based mechanism reconciles prior observations and provides a foundation for designing next-generation peptide inhibitors.

## MATERIALS AND METHODS

### Protein expression and purification

NDM-1 (residues G29–R270; Uniprot C7C422) was expressed from pET28a(+) TEV vector in *E. coli* BL21 cells. Isotopically labeled samples (¹⁵N-and ¹³C/¹⁵N) were prepared using a high-density M9 minimal medium shift protocol. Protein expression was induced with 1 mM IPTG at 25 °C for 20 h. His_6_-NDM-1 was purified by Ni-affinity chromatography, cleaved with TEV protease (1:100 protein:TEV molar ratio), and a second Ni-affinity step was used to remove the His₆ tag and His-tagged TEV (cleaved NDM-1 collected in the flow-through). The protein was buffer-exchanged into 100 mM Bis-Tris-HCl, 150 mM NaCl, pH 7.0 and stored at −80 °C. Full experimental details are provided in the Supporting Information.

### NMR studies

All NMR experiments used tag-cleaved NDM-1 prepared in the holo state by addition of 4 equivalents ZnCl₂ per monomer; samples were exchanged into phosphate-free 100 mM Bis-Tris-HCl, 150 mM NaCl, pH 7.0 (5–10% D₂O) to maintain zinc solubility. Spectra were recorded at 700 MHz on a cryogenically cooled probe at 30 °C. Backbone assignments were taken from deposited BMRB entries and verified with standard TROSY-based triple-resonance experiments. Chemical-shift perturbations were measured from^1^H–^15^N TROSY-HSQC titrations with thanatin; backbone dynamics were assessed by TROSY R_1_ and R_2_ measurements. Intermolecular NOEs were identified using^15^N/^13^C-filtered NOESY-HSQC experiments and cross-checked against peptide-only and apo controls. Full acquisition parameters, titration conditions, relaxation delays, and processing details are provided in the Supporting Information.

### Molecular docking

Thanatin–NDM-1 docking was performed using HADDOCK 2.4^12,13^, guided by NMR chemical shift perturbations and intermolecular NOEs to define active residues on NDM-1. Thanatin was modeled from its solution NMR structure, and docking proceeded through rigid-body, semi-flexible, and water-refinement stages. Histidine protonation states were explicitly defined to reflect experimental pH 7. The top-scoring cluster was selected for analysis, and full docking parameters and scoring statistics are provided in the Supporting Information.

### Molecular dynamics

MD simulations were carried out with GROMACS using the AMBER99SB-ILDN force field^14,15^ to examine the structural dynamics of NDM-1 in its apo and thanatin-bound states. Three independent 150 ns simulations were first performed to characterize global conformational behavior and identify representative peptide-bound configurations. To resolve loop dynamics within these distinct binding states, five 20 ns simulations were performed for each state (apo and peptide-bound). All systems were simulated in explicit solvent under standard NVT/NPT conditions. To preserve the experimentally observed Zn²⁺ coordination geometry, harmonic distance restraints were applied between the zinc ions and their coordinating residues during equilibration and production. These restraints did not involve residues at the thanatin-binding interface. Analyses focused on RMSD and RMSF to assess peptide-dependent changes in protein flexibility. Full simulation parameters and input files are provided in the Supporting Information.

### Circular Dichroism (CD) Thermal Denaturation

Thermal denaturation of NDM-1 was monitored by circular dichroism at 222 nm using a 1 mm pathlength quartz cuvette. Protein samples (35 μM) were heated from 20 to 85 °C at 1 °C/min. Data were normalized and fitted to a Boltzmann sigmoidal model to extract apparent melting temperatures (T_m_). Reported values represent mean ± SD from three independent replicates. Additional experimental details, raw traces, fit parameters, and transition widths (dT) are provided in the Supporting Information.

### Synergy assay

*E. coli* BL21 expressing NDM-1 from pET28a(+)-TEV, a non-expressing wild-type BL21 control, and a TEV-expressing control were grown overnight in LB with appropriate antibiotics, subcultured into fresh LB containing 1 mM IPTG, and grown 4–5 h before normalization to OD_600_ = 1.0. Cultures were diluted 1:100 into 96-well plates containing LB, LB + thanatin (6 μg·mL⁻¹), or LB + thanatin (6 μg·mL⁻¹) + imipenem (1 μg·mL⁻¹); all wells contained 1 mM IPTG. Plates were incubated at 37 °C with continuous orbital shaking and OD_600_ recorded every 15 min for 8 h. Input and output viable counts were determined by serial dilution and spot plating; data were analyzed in GraphPad Prism 9 (GraphPad Software, San Diego, CA, USA). Full experimental details (strain genotypes and plasmids, antibiotic sources and concentrations, exact inoculation volumes, plate-reader model and settings, number of biological and technical replicates, plating volumes, and statistical tests) are provided in the Supporting Information.

## RESULTS AND DISCUSSION

### Thanatin Engages the Catalytic Groove of NDM-1 through a Surface-Exposed β-Hairpin Interface

To identify residues involved in thanatin recognition, solution-state NMR spectroscopy was used to map chemical-shift perturbations (CSPs) and intermolecular contacts. Two-dimensional^1^H–^15^N HSQC spectra of holo-NDM-1 acquired during thanatin titration confirmed preservation of the global protein fold (Figure 1A). Significant CSPs (> 0.017 ppm) defined a continuous interaction surface encompassing residues Phe21–Asp23, Leu40–Gly46, Val48, Trp68–Thr73, Trp79, His97–Asp99, Gly102–Leu107, and Asn117–Ala131 (Figure 1B–E). Confirmation that peptide binding does not induce zinc release comes from the comparison of apo– and holo-NDM-1 spectra, which reveals distinct metal-dependent chemical-shift patterns (Figure S1A). During thanatin titration, NDM-1 retained spectral features characteristic of the holo-state, whereas subsequent EDTA treatment reproduced the apo-fingerprint (Figure S1B).

**Figure 1.**
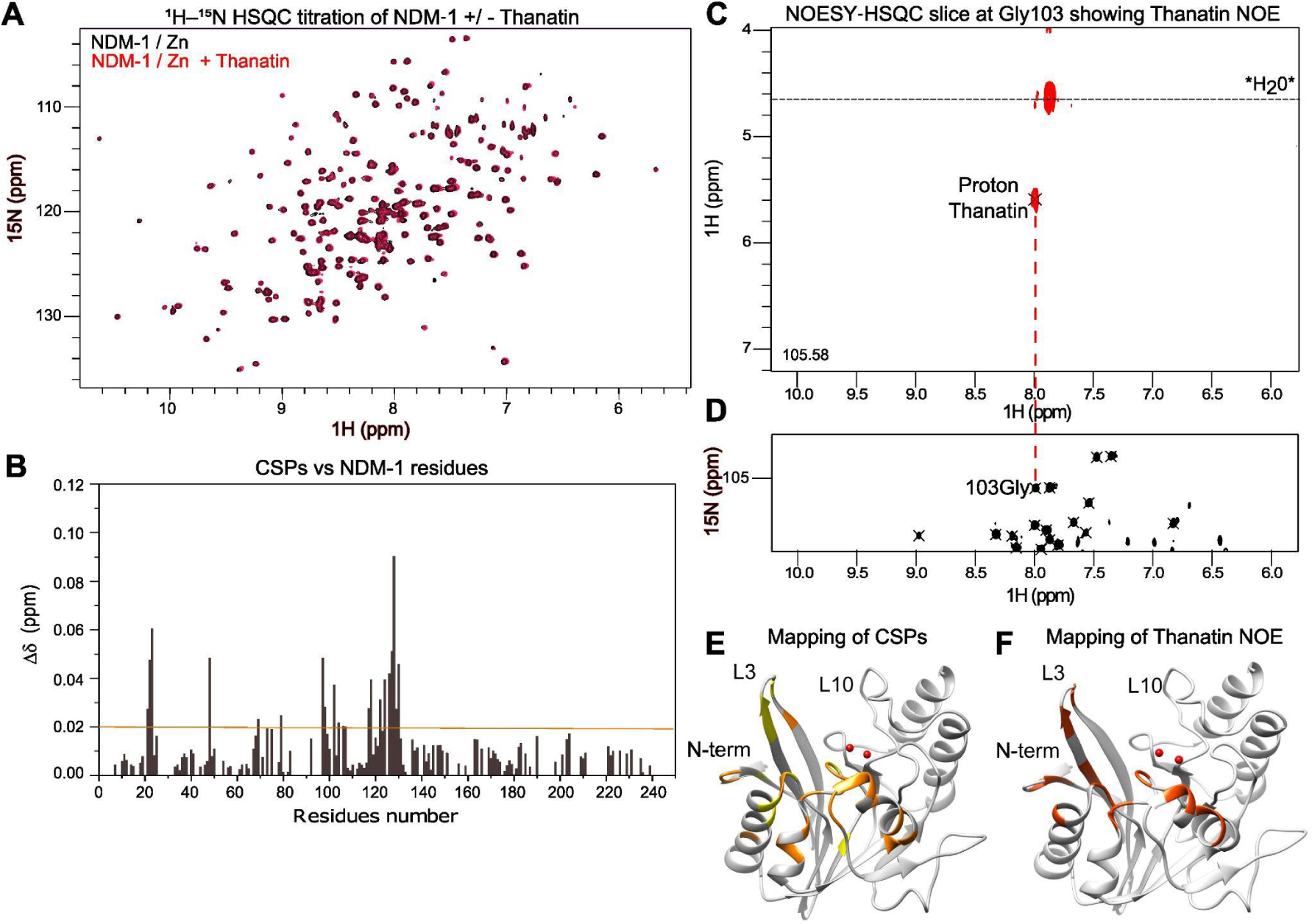
Thanatin Binding Induces Chemical Shift Perturbations and a Specific Intermolecular Contact in NDM-1. (A) Overlay of ¹H–¹⁵N HSQC spectra of Zn–NDM-1 recorded in the absence (black) and presence (red) of thanatin, showing widespread chemical shift perturbations upon peptide binding. (B) Per-residue weighted ¹H–¹⁵N chemical shift perturbations (Δδ) of Zn–NDM-1 induced by thanatin. The horizontal dashed line indicates the threshold used to identify significantly perturbed residues. (C) Representative 3D ¹H–¹H–¹⁵N NOESY–HSQC plane extracted at the ¹⁵N frequency of Gly103 (105.58 ppm), revealing an intermolecular NOE between the Gly103 amide proton and a thanatin proton; the residual H₂O signal is indicated. (D) ¹H–¹⁵N HSQC spectrum of Zn–NDM-1 highlighting Gly103; a dashed line links the selected amide resonance to the corresponding NOESY plane shown in panel C, confirming residue-specific assignment of the intermolecular NOE. (E) Mapping of residues exhibiting significant CSPs (panel B) onto the crystal structure of NDM-1 (PDB: 6TTC). Residues with large CSPs (Δδ > 0.02 ppm) are shown in orange, and those with moderate but significant CSPs (0.015 ppm < Δδ ≤ 0.02 ppm) are shown in yellow, highlighting perturbed regions in the N-terminus and loops L3 and L10. (F) Structural mapping of all observed intermolecular NOEs between NDM-1 and thanatin (red) onto the same structure, demonstrating that the NOE-defined contacts fall within the CSP-defined binding interface.

^15^N/^13^C-filtered NOESY-HSQC experiments identified 43 intermolecular NOEs between thanatin side-chain protons and NDM-1 amide resonances (Figure 1C–D; Table S1). These contacts mapped to the N-terminal segment (Gln13, Met14, Phe21), β3/β4 sheets (Thr37, Phe45), helix α1 (Trp68, Trp79), the zinc-proximal Asp99, and helix α3 (Gln122, Leu123) (Figure 1F). The NOE distribution closely mirrored the CSP-defined interface, confirming a contiguous binding surface adjacent to the catalytic groove.

Together, these data show that thanatin engages a peripheral surface of NDM-1 while preserving its native zinc-bound state under our solution-state NMR conditions. This outcome contrasts with the zinc-displacement model proposed by Ma et al.^5^ and supports a zinc-retaining allosteric mechanism.

### NMR-Guided HADDOCK Docking Defines an Allosteric Binding Mode

To generate a structural model of the complex, HADDOCK docking was performed using experimental restraints derived from CSPs (> 0.017 ppm) and intermolecular NOEs. Thanatin was modeled from its solution NMR structure (PDB 00006AAB), which adopts a disulfide-stabilized β-hairpin fold.

In the lowest-energy models, thanatin binds a surface region adjacent to the L3 loop, with its cationic N-terminus oriented toward a negatively charged patch while remaining distal from the catalytic dinuclear zinc center (Figure 2A). Multiple clusters were examined, and the top-scoring cluster (HADDOCK score = −95.5 ± 8.2; 11 structures; RMSD ≈ 1 Å) exhibited favorable interaction energies and the largest buried surface area (724 Å²; Table S3). The interface involves salt bridges, hydrogen bonds, and hydrophobic contacts engaging the N-terminal extension, β3/β4 sheet, loop L3, and helices α1–α3 (Figure 2B; Table S4).

**Figure 2.**
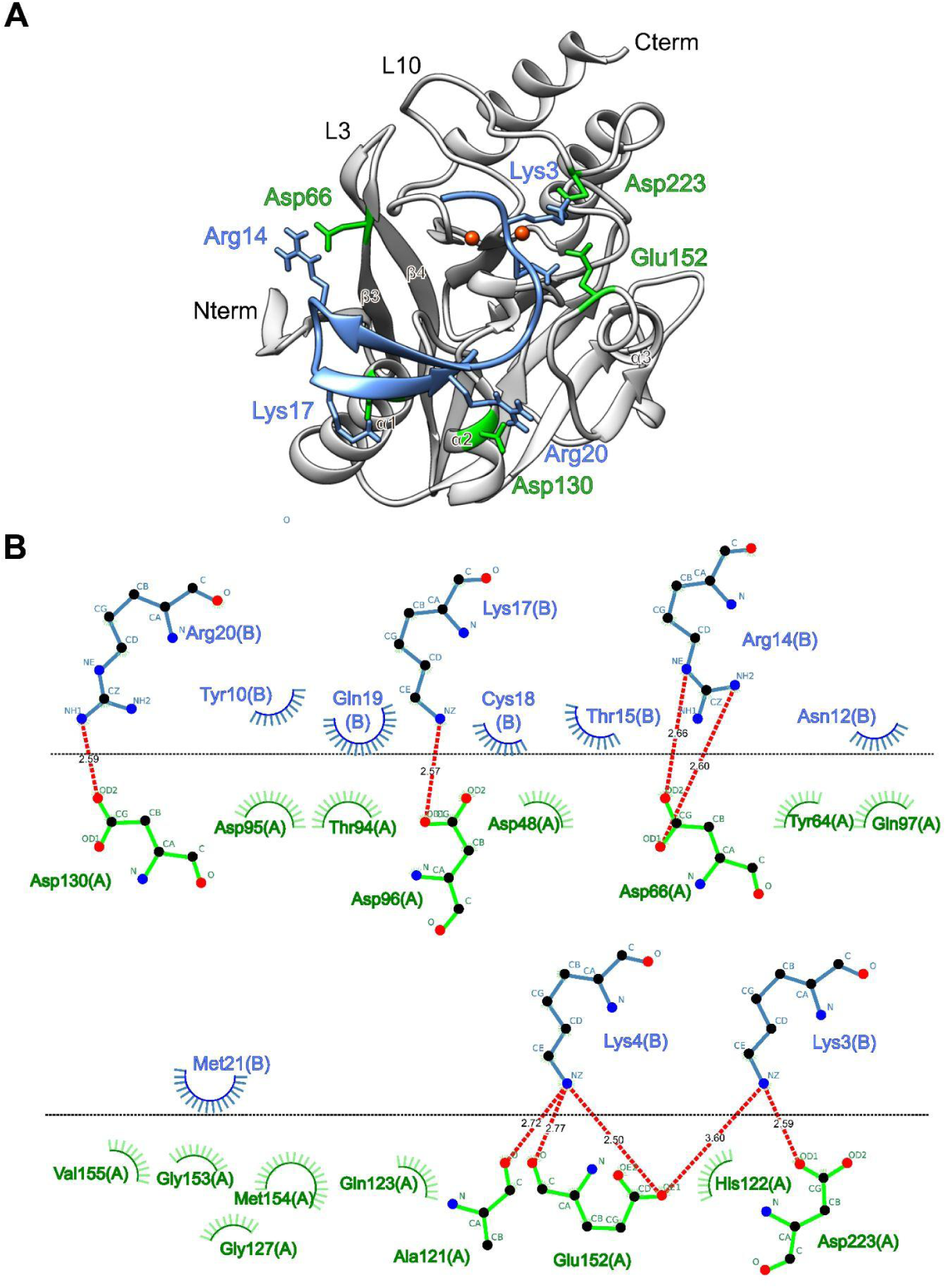
NMR-guided docking model of the NDM-1–thanatin complex. (A) Representative low-energy HADDOCK model of thanatin bound to NDM-1 generated using NMR-derived restraints. The NDM-1 structure (PDB: 5ZR8) is shown in cartoon representation (grey), with residues identified by NMR chemical shift perturbations and intermolecular NOEs highlighted in green. Thanatin is shown in teal. The model positions thanatin in proximity to the L3 loop of NDM-1, consistent with the NMR-defined binding surface shown in Figure 1.(B) Predicted interaction map derived from the docking model, generated using LigPlot+. Residues involved in putative intermolecular contacts are displayed in ball-and-stick representation, and predicted hydrogen bonds are indicated by red dashed lines with interatomic distances (Å). This analysis illustrates the types of interactions that may contribute to stabilization of the complex within the proposed docking model. Complete residue-level interaction data are provided in SI Table 2.

Importantly, the HADDOCK output represents a restraint-guided ensemble rather than a definitive structure, and full validation of the interface will require complementary experiments such as targeted mutagenesis or additional independent restraints. Nonetheless, the modeled binding site is spatially distinct from the catalytic zinc center and flexible loop L10, supporting an allosteric binding mode rather than metal chelation. While this docking ensemble suggests a peripheral interface, we next sought to experimentally verify that this surface interaction does not destabilize the enzyme’s core or displace the catalytic metal ions.

### Thanatin Binding Preserves the Catalytic Di-zinc Architecture

To evaluate whether inhibition involves metal displacement, we monitored NDM-1 thermal denaturation by circular dichroism (CD) at 222 nm (Figure 3). Coordination of Zn²⁺ to apo-NDM-1 (T_m_ = 49.5 ± 0.03 °C) markedly stabilized the protein (holo-NDM-1; Tm = 58.2 ± 0.1 °C). Addition of thanatin to the holo-enzyme produced a T_m_ (57.5 ± 1.1 °C, n = 3, p = 0.29) indistinguishable from the zinc-bound state. CD cannot distinguish unfolding from aggregation, a limitation that is especially relevant for the apo state.

**Figure 3.**
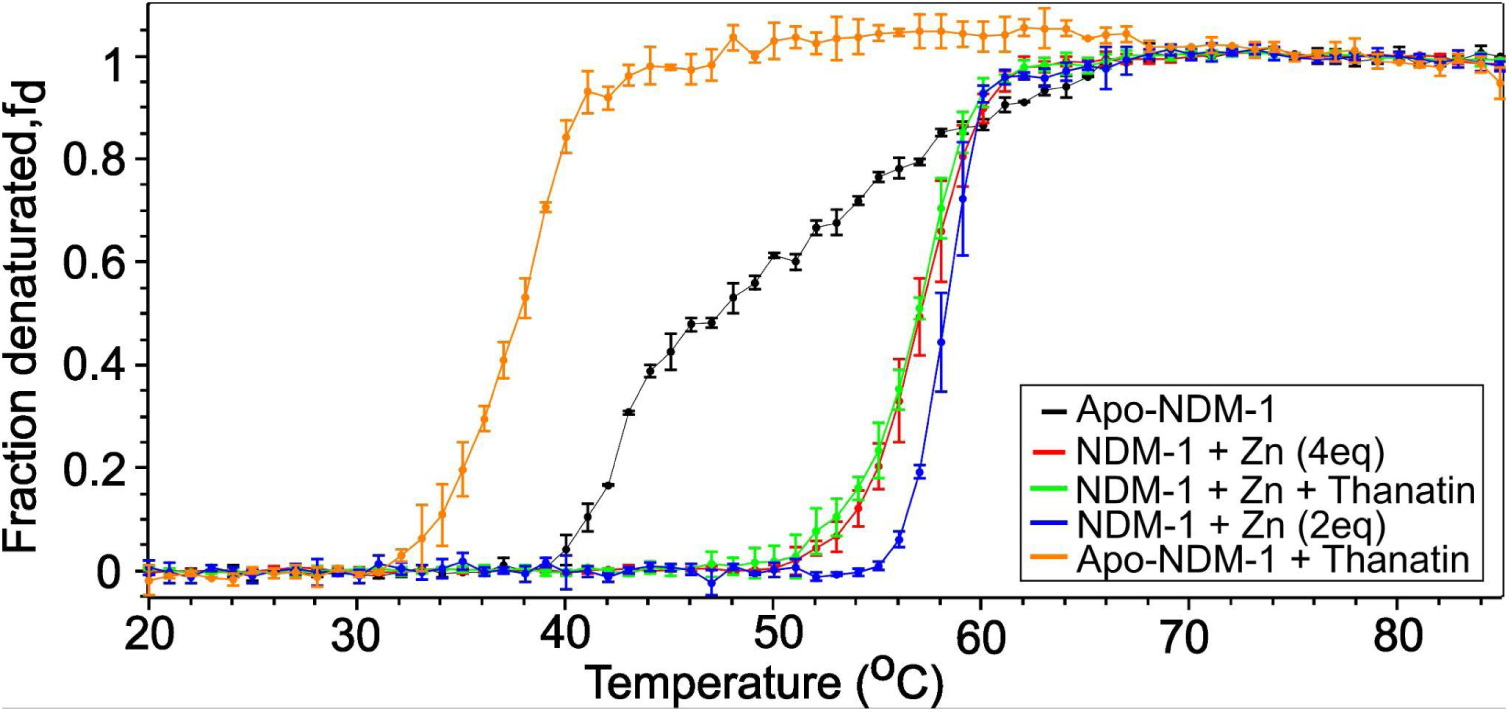
CD thermal denaturation reveals the metal-dependent stability and peptide-binding landscape of NDM-1. CD unfolding curves for NDM-1 under five conditions (apo-NMD-1, apo-NDM-1 + thanatin, Zn-bound at 2 and 4 eq, and Zn + thanatin). Apo NDM-1 unfolds broadly (T_m_ = 49.5 ± 0.03 °C), whereas thanatin binding sharpens the transition but lowers stability (T_m_ = 37.8 ± 0.06 °C), consistent with Zn-independent association that alters cooperativity. Zn²⁺ binding stabilizes the fold (T_m_ = 58.2 ± 0.07 °C at 2 eq; 57.1 ± 0.15 °C at 4 eq), and the Zn + thanatin curve (T_m_ = 56.8 ± 0.04 °C) overlaps Zn alone, indicating that thanatin neither displaces Zn nor further stabilizes the holo-enzyme. Curves represent mean ± SD (n = 3). Fits were obtained using a Boltzmann sigmoidal model; full parameters are listed in Table SI2.

Even so, the overlapping Tm values show that the zinc-bound scaffold maintains comparable global stability in the presence of thanatin. This similarity reflects overall stability rather than identical conformational microstates, which CD cannot resolve. This thermal behavior, together with preservation of the NMR zinc fingerprint (Figure 1A), rules out a zinc-release mechanism.

In contrast, the transition from holo– to apo-NDM-1 is accompanied by a dramatic loss of stability (ΔT_m_ ≈ 9 °C). If thanatin removed zinc, NDM-1 would be expected to show similar destabilization. Instead, the protein remains folded, indicating that inhibition arises from allosteric interactions within the native architecture rather than metal removal.

Interestingly, thanatin markedly destabilized apo-NDM-1 (T_m_ = 37.8 ± 0.06 °C; p < 0.0001). This shift reflects altered folding cooperativity rather than complete global unfolding. It indicates that thanatin can engage the apo scaffold, but that the di-zinc center is required for the peptide to stabilize a productive inhibitory conformation (Figure SI2, Table SI2). Having confirmed that the catalytic center remains intact within the thanatin–NDM-1 complex, we employed Molecular Dynamics (MD) simulations to investigate how this distal binding modulates the essential protein fluctuations required for turnover.

### Mechanistic Basis of Inhibition: Allosteric Rigidification of the L3 Loop

To define how thanatin binding perturbs the functional dynamics of NDM-1, we employed molecular dynamics (MD) simulations to interrogate conformational sampling in both apo and peptide-bound states. Triplicate extended simulations (150 ns each) confirmed that the catalytic di-zinc center remains structurally rigid throughout (Figure SI3), consistent with NMR and CD measurements indicating preserved metal coordination. In contrast, thanatin binding exerted a localized effect on the intrinsic dynamics of the L3 loop, a flexible structural element essential for substrate access and catalysis.

Quantitative analysis of L3-loop conformational variability revealed a clear reduction in ensemble breadth upon peptide binding. When RMSD values were calculated for the L3 loop following alignment on the protein core (excluding loop residues), the apo enzyme displayed broad, high-amplitude sampling (RMSD = 0.247 ± 0.075 nm), whereas the thanatin-bound state populated a markedly narrower ensemble (RMSD = 0.186 ± 0.049 nm; Figure 4A,C). These data indicate that peptide engagement restricts L3-loop mobility without imposing a single static conformation, consistent with an ensemble-based mode of inhibition.

**Figure 4.**
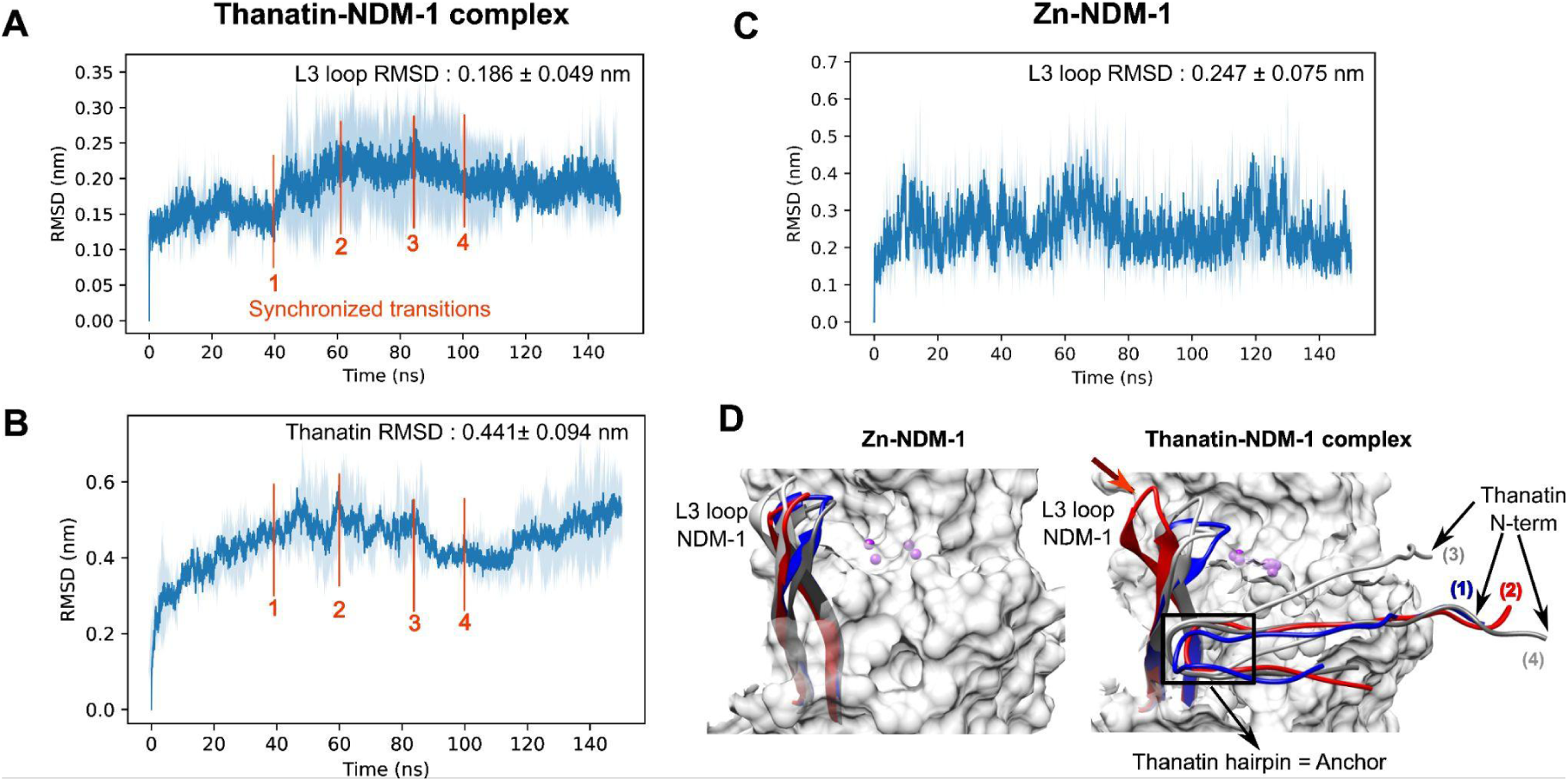
Conformational sampling and dynamic coupling of the thanatin–NDM-1 complex. (A) RMSD profile of the NDM-1 L3 loop in the thanatin-bound state (0.186 ± 0.049 nm). (B) RMSD profile of thanatin (0.441 ± 0.094 nm). Red vertical markers (1–4) denote synchronized conformational transitions at ∼40, ∼60, ∼85, and ∼100 ns. Across the trajectory, shifts in peptide RMSD coincide with L3-loop reorganizations, indicating strong dynamic coupling in which the loop adjusts to maintain an optimized interface as thanatin samples metastable poses. (C) Baseline RMSD profile of the apo L3 loop (0.247 ± 0.075 nm), showing higher-amplitude sampling and the absence of the restricted, synchronized fluctuations observed in the bound complex. Shaded regions represent SD across triplicate 150 ns simulations. (D) Structural snapshots of the apo enzyme (left) and the thanatin–NDM-1 complex (right) at 20 ns (blue), 40 ns (red), 60 ns (dark grey), and 100 ns (light grey). At 40 ns, the complex undergoes a transient excursion in which the N-terminus and L3 loop lift (marked by the red arrow) from the active site, before the hairpin turn facilitates a rapid return to the inhibitory pose by 60 ns. Zinc coordination remained stable throughout all trajectories (Figure SI3).

Analysis of peptide dynamics shows that thanatin remains confined near the NDM-1 L3 loop while sampling multiple metastable binding poses. Peptide RMSD fluctuations are temporally coupled to subtle reorganizations of the L3 loop, reflecting correlated motions between binding partners (Figure 4A,B). At ∼40 ns, the loop transiently lifts away from the catalytic site, accompanied by local rearrangements of the thanatin N-terminus and hairpin, while the peptide remains peripherally bound. Recovery to the inhibitory pose occurs by ∼60 ns, facilitated by the central hairpin and C-terminus. Together, these observations show that the peptide remains peripherally associated even during occasional large-amplitude excursions of the L3 loop (Figure 4B,D).

To rigorously quantify loop dynamics across binding states, five independent 20 ns simulations were initiated from representative configurations. These shorter trajectories were used as ensemble sampling rather than full convergence, enabling assessment of reproducible trends across independently seeded simulations. Apo-NDM-1 exhibited elevated backbone RMSF values throughout the L3 region, reflecting substantial intrinsic flexibility (Figure 5B). Thanatin binding selectively attenuated these fluctuations, yielding a statistically significant reduction in mean L3-loop RMSD relative to the apo enzyme (Apo: 0.159 ± 0.007 nm; Bound: 0.142 ± 0.006 nm; n = 5; Welch’s t-test, p = 0.0043; Figure 5C). Importantly, this effect was spatially confined to the peptide-proximal interface; the distal L10 loop (residues 210–230) exhibited increased flexibility in the bound state (Figure 5B), indicating redistribution rather than global suppression of protein dynamics.

**Figure 5.**
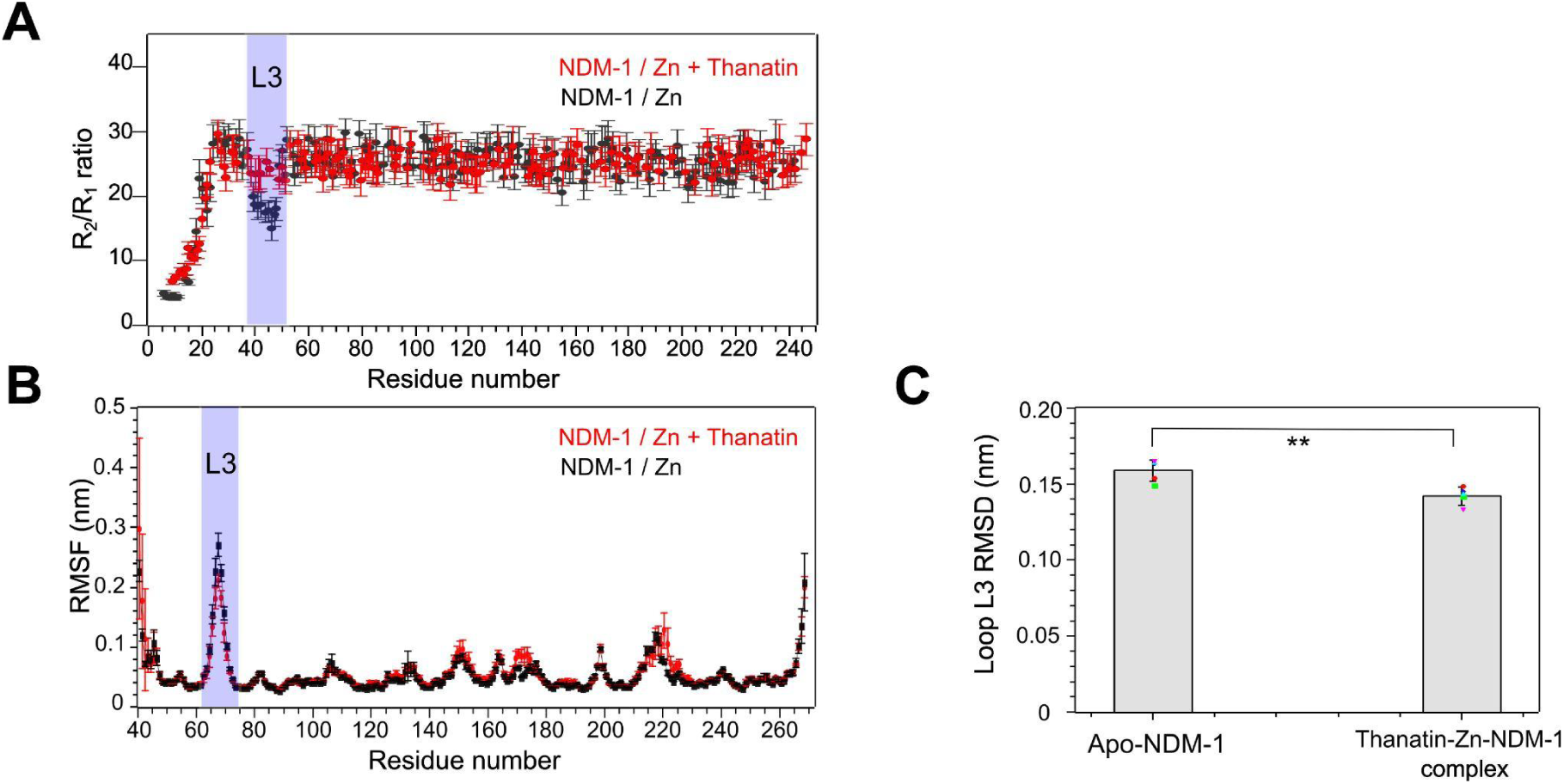
Thanatin restricts the conformational ensemble of the NDM-1 L3 loop. (A) ¹⁵N NMR R₂/R₁ relaxation ratios for apo-NDM-1 (black) and the thanatin-bound complex (red) plotted against residue number. The blue shaded region highlights elevated R₂/R₁ ratios within the L3 loop upon binding, indicating localized rigidification on the ps–ns timescale. (B) Root-mean-square fluctuation (RMSF) profiles from five independent 20 ns MD replicates (n = 5) for apo-NDM-1 (black) and the thanatin-bound complex (red). Simulations were initiated from the NDM-1 crystal structure (PDB ID: 00003ZR9, chain A), with residue numbering beginning at position 40. Thanatin binding markedly reduces fluctuations in the L3 loop (blue shading), whereas the distal L10 loop (residues 210–230) retains high mobility. (C) Statistical comparison of L3-loop backbone RMSD across replicates. Data are shown as mean ± SD with individual replicate values overlaid (Apo: 0.159 ± 0.007 nm; Complex: 0.142 ± 0.006 nm). Welch’s two-tailed t-test indicates a significant reduction in L3-loop mobility (**p = 0.0043).

Experimental NMR relaxation measurements independently corroborated this localized anchoring mechanism. Residues within the L3 loop exhibited elevated R₂/R₁ ratios upon thanatin binding (Figure 5A), consistent with attenuation of fast (ps–ns) backbone motions. The absence of intermolecular tr-NOE signals (Figure 1F) involving L10 residues further confirms that increased mobility in this distal loop arises from indirect dynamic compensation rather than direct peptide engagement.

Collectively, these results establish a zinc-retaining dynamic allosteric mechanism in which thanatin limits L3-loop flexibility while leaving metal coordination and the global fold intact (Figures 3 and SI6). This illustrates how antimicrobial peptides can exploit intrinsic loop dynamics to inhibit the enzyme through conformational restriction.

### Dynamic Allosteric Restriction of the L3 Loop Drives Synergistic Restoration of Imipenem Efficacy in NDM-1 Pathogens

To bridge our structural observations with biological function, we challenged NDM-1-expressing pathogens with a combination of thanatin and imipenem, a synthetic β-lactam antibiotic that is efficiently hydrolysed by NDM-1. Wild-type and empty-vector controls confirmed that the observed effects were specifically NDM-1 mediated (Figure SI4). While thanatin remains non-toxic at 6 μg/mL (Figure 6A), it partially restores imipenem susceptibility, as reflected by the immediate suppression of growth upon antibiotic addition (Figures 6B–E).

**Figure 6.**
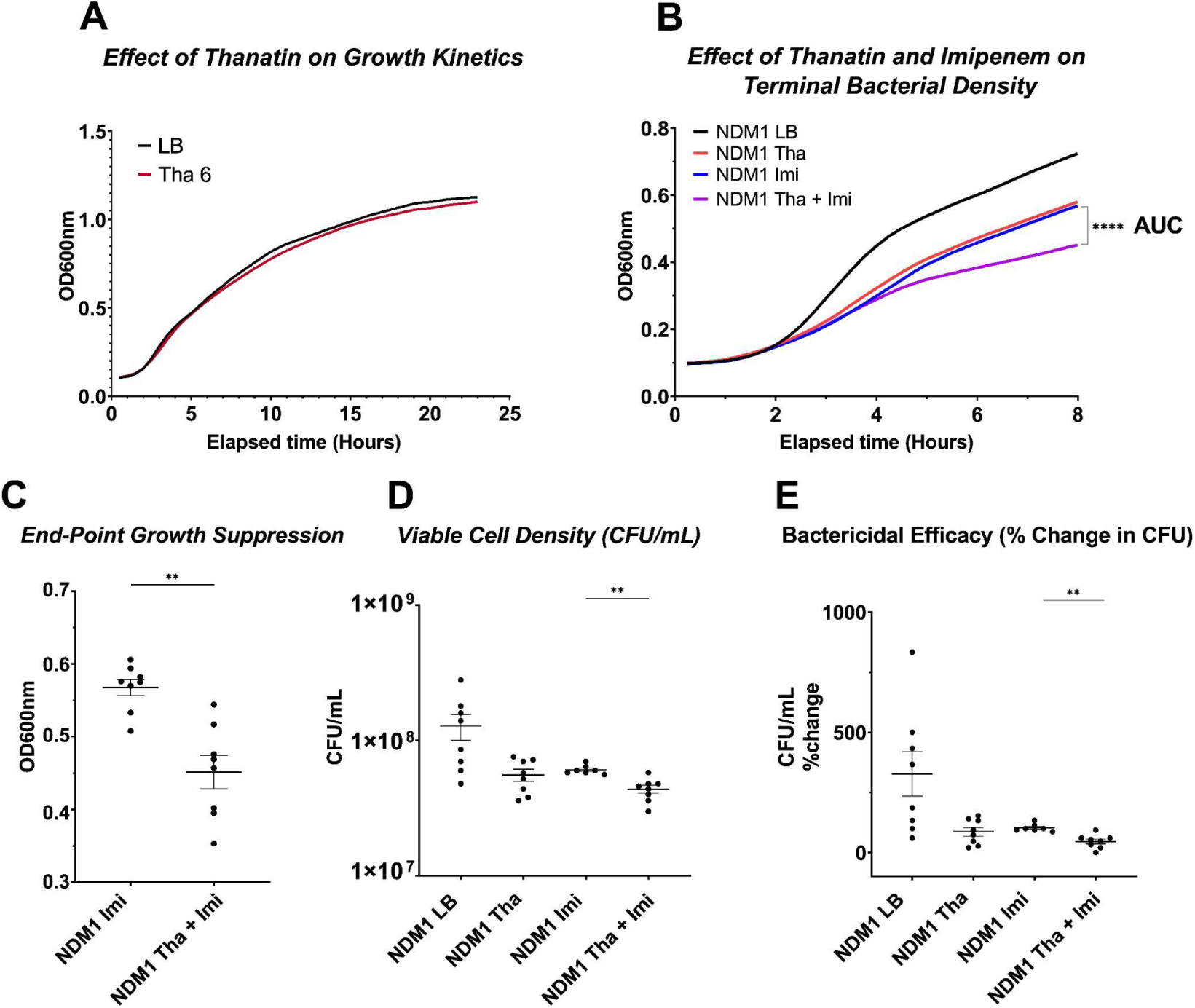
Thanatin synergizes with imipenem through dynamic allosteric inhibition of NDM-1. (A) Growth kinetics of *E. coli* BL21 expressing NDM-1 in the presence or absence of thanatin (6 µg/mL). (B) Growth dynamics over eight hours with imipenem (1 µg/mL) alone or in combination with thanatin. Lines represent the mean of n = 8 replicates. Statistical significance was assessed by AUC analysis using an unpaired t-test (p < 0.0001). (C) Terminal OD_600_ values at eight hours, showing a 19.5% reduction in growth for the combination treatment relative to imipenem alone. (D) Viable cell density (CFU/mL) and (E) percent change in CFU/mL relative to initial inoculum for the indicated treatments. Individual points represent independent replicates (n = 7–8); one outlier in the imipenem condition was removed using the ROUT method (q = 1%). Horizontal lines and error bars denote mean ± SEM. Statistical significance in panels C–E was determined using a two-tailed Mann–Whitney U test (ns, * p < 0.05, ** p < 0.01, *** p < 0.001, **** p < 0.0001). The observed phenotypic synergy at 37 °C—manifested as suppressed growth and reduced viability—is consistent with the high thermal stability of the thanatin–NDM-1 complex (T_m_ = 57.5 °C vs. 37.8 °C for the apo form) and supports a dynamic allosteric mechanism in which thanatin rigidifies the L3 loop while preserving the catalytic di-zinc center.

Because thanatin does not act as a classical competitive inhibitor of NDM-1, functional synergy (UAI) provides the most relevant measure of the biological consequence of peptide binding. Synergy assays were performed under high IPTG induction of NDM-1, resulting in strong overexpression of the enzyme. Under these stringent conditions, the observed sensitization underestimates the effect that would occur at physiological NDM-1 levels.

To quantify this effect, inhibitory efficacy was assessed by comparing growth dynamics up to the onset of stationary phase (8 h), a window chosen to avoid artificial inflation of AUC values from prolonged stationary-phase measurements. Statistical analysis revealed that the thanatin–imipenem combination significantly suppressed the NDM-1 strain compared to the imipenem-only control (AUC = 2.150 ± 0.045 vs. 2.425 ± 0.029; Figure 6B). This suppression was further reflected by a 19.5% reduction in terminal OD600 (p = 0.0011; Figure 6C) and corroborated by viable cell counts (CFU/mL). While imipenem alone permitted a 103% expansion in viable cells, the combination restricted this expansion to 46% – a modest but reproducible sensitization consistent with our proposed dynamic allosteric mechanism (p = 0.0011; Figures 6D–E).

These phenotypic results support our zinc-retaining dynamic allosteric model over previously proposed zinc-release mechanisms^5^. While it has been suggested that thanatin triggers zinc displacement, our CD analysis (Figure 3) demonstrates that the thanatin–NDM-1 complex is highly stable (T_m_ = 57.5 °C), whereas the apo-form is structurally compromised at physiological temperatures (T_m_ = 37.8 °C). This thermal instability implies that a zinc-release mechanism would result in structural collapse at 37 °C, precluding the consistent synergy observed in our assays. Instead, our NMR and EDTA-titration data confirm that the di-zinc center remains intact, while MD simulations corroborate a reduction in the conformational plasticity of the L3 loop.

## CONCLUSION

The New Delhi Metallo-β-lactamase (NDM-1) is a major driver of global multidrug resistance, neutralizing even “last-resort” carbapenems. Here, we define how the antimicrobial peptide thanatin inhibits this enzyme. Our results reveal an allosteric mechanism that may be exploited to restore the efficacy of existing antibiotics.

The catalytic activity of NDM-1 is fundamentally dependent on its di-zinc active center, which serves to activate the nucleophilic water molecule required for β-lactam hydrolysis^16,17^. Previously, Ma et al.^5^ proposed that the antimicrobial peptide thanatin inhibits NDM-1 by inducing the release of these essential zinc ions.^5^ However, our results reveal a distinct mode of action. High-resolution NMR titrations and EDTA-challenge experiments demonstrate that the di-zinc coordination remains intact upon peptide engagement. These observations contrast with the earlier metal displacement hypothesis^5^ and instead indicate that, under native-like conditions, thanatin inhibits NDM-1 while the catalytic center remains fully coordinated, preserving the holo-enzyme state. By binding to a site adjacent to the catalytic groove, thanatin restricts the motion of the inherently flexible L3 loop. NMR relaxation measurement and MD simulations both reveal localized rigidification of this region, supporting a model in which thanatin limits the conformational fluctuations required for efficient substrate turnover. This perspective points toward a shift in strategy for metallo-β-lactamase inhibition: from competitive metal sequestration to the targeted modulation of local protein dynamics.

This mechanistic insight carries clinical relevance given the increasing prevalence of carbapenem-resistant *E. coli* strains that compromise the efficacy of broad-spectrum antibiotics such as imipenem^18,19^. While thanatin has been explored for its intrinsic antimicrobial activity^20,21^, our findings suggest that it may also function as an adjuvant capable of enhancing existing therapies. Current clinical combinations, such as imipenem with the β-lactamase inhibitor relebactam^22,23^, remain largely ineffective against NDM-type enzymes^22,24^, underscoring the need for alternative approaches.

Our synergy experiments show that thanatin increases the susceptibility of NDM-1–producing pathogens at clinically relevant antibiotic concentrations. Combining subinhibitory thanatin with imipenem reduced bacterial viability relative to either monotherapy, and this effect remained detectable even under high IPTG-induced NDM-1 expression. Because this induction level exceeds typical clinical expression, the observed sensitization likely represents a conservative estimate of the peptide’s functional impact. Clinically prevalent variants such as NDM-5 and NDM-7 carry substitutions distal to the thanatin-binding surface. This structural conservation suggests that the allosteric mechanism identified here may extend to these alleles; this possibility now merits direct experimental testing.

Ultimately, our findings redefine the mechanism of thanatin-mediated NDM-1 inhibition. Rather than promoting loss of the catalytic zinc ions, thanatin preserves a fully coordinated di-zinc active center while allosterically restricting the L3 loop. This localized rigidification limits the conformational fluctuations and stabilizes the enzyme in a less active, yet folded, state. More broadly, these results identify protein dynamics – rather than metal sequestration – as a tractable vulnerability in NDM-1. Exploiting this allosteric, dynamics-based mechanism may provide a foundation for next-generation antibiotic adjuvants that restore carbapenem efficacy against metallo-β-lactamase-producing pathogens.

## SUPPORTING INFORMATION

Supporting Information is available and includes Supplementary Figures S1–S6, Supplementary Tables S1–S4, extended NMR experimental methods, full CSP and intermolecular NOE datasets, HADDOCK docking parameters and scoring statistics, molecular dynamics simulation details (including zinc-coordination RMSD, RMSF profiles, and replicate trajectories), CD thermal-denaturation fitting parameters and replicate traces, and additional bacterial growth controls and synergy assay methods. This material is available free of charge via the Internet at http://pubs.acs.org.

## AUTHOR CONTRIBUTIONS

GR and LJM designed the study. PK performed and analyzed the molecular docking and molecular dynamics simulations, and prepared the corresponding sections of the manuscript. GR designed and carried out the laboratory experiments, data analysis, and data integration. T.C. and A.H. conducted the microbiological experiments, analyzed the associated data, and wrote the corresponding sections. GR and LJM prepared and edited the manuscript with input from all authors.

## NOTES

The authors declare no competing financial interest.

## ACKNOWLEDGMENT

Research reported in this publication was supported by the National Institutes of Health under award number GM145369 to L.J.M. and awards R35GM158026, R01AI157106, R01AI178908, R01AI181382, and R21AI181381 to A.H. Computations were performed using the computer clusters and data storage resources of the HPCC, which were funded by grants from NSF (MRI-2215705, MRI-1429826) and NIH (1S10OD016290-01A1).

## ABBREVIATIONS

AMR: antimicrobial resistance
CD: circular dichroism
CSP: chemical-shift perturbation
EDTA: ethylenediaminetetraacetic acid
HADDOCK: high ambiguity driven protein-protein DOCKing
HSQC: heteronuclear single quantum coherence
IPTG: isopropyl β-D-1-thiogalactopyranoside
LB: Luria-Bertani broth
MD: molecular dynamics
MDR: multidrug-resistant
NDM-1: New Delhi metallo-β-lactamase 1
NMR: nuclear magnetic resonance
NOE: nuclear Overhauser effect
NOESY: nuclear Overhauser effect spectroscopy
OD_600_: optical density at 600 nm
PDB: Protein Data Bank
RMSD: root-mean-square deviation
RMSF: root-mean-square fluctuation
SD: standard deviation
TEV: tobacco etch virus
T⍰: apparent melting temperature
TROSY: transverse relaxation-optimized spectroscopy

## SYNOPSIS

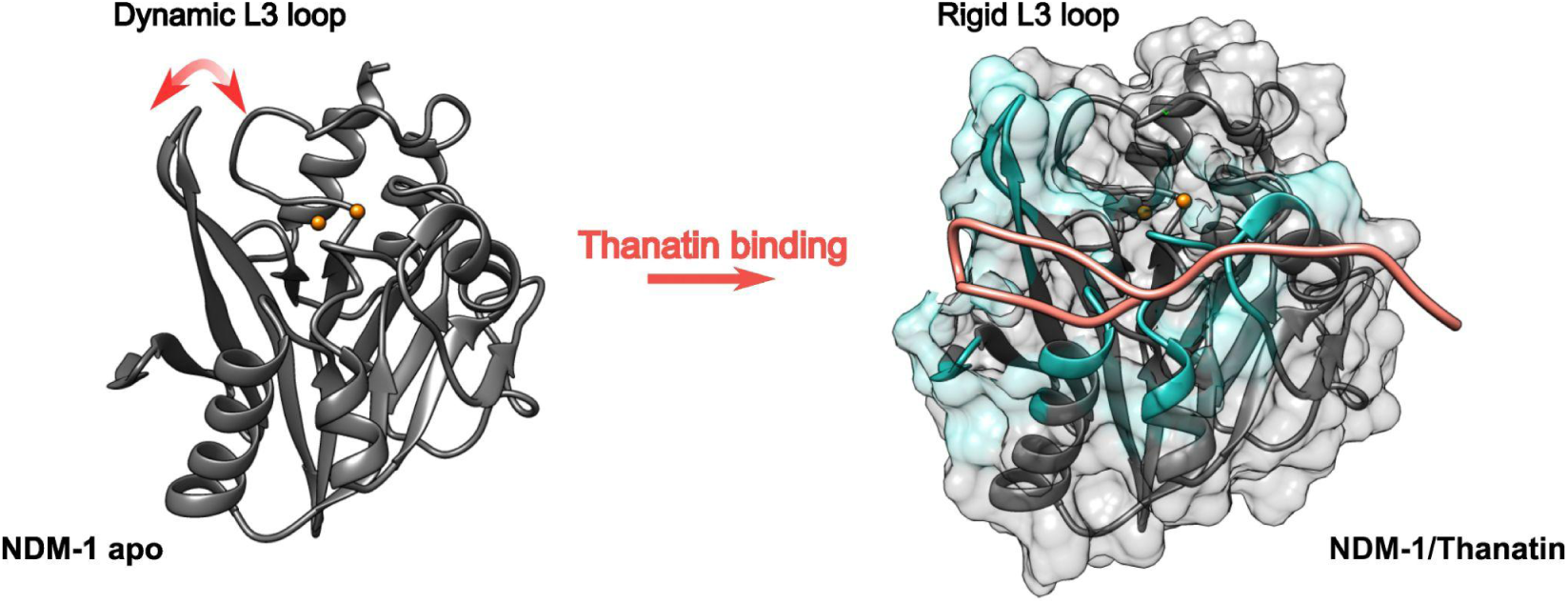
Thanatin binds adjacent to the catalytic groove, rigidifying the NDM-1 L3 loop while preserving the native di-zinc center.

## REFERENCE.

(1) Antimicrobial Resistance Collaborators. Global Burden of Bacterial Antimicrobial Resistance in 2019: A Systematic Analysis. Lancet Lond. Engl. 2022, 399 (10325), 629–655. 10.1016/S0140-6736(21)02724-0.

(2) Macesic, N.; Uhlemann, A.-C.; Peleg, A. Y. Multidrug-Resistant Gram-Negative Bacterial Infections. The Lancet 2025, 405 (10474), 257–272. 10.1016/S0140-6736(24)02081-6.

(3) Feng, H.; Liu, X.; Wang, S.; Fleming, J.; Wang, D.-C.; Liu, W. The Mechanism of NDM-1-Catalyzed Carbapenem Hydrolysis Is Distinct from That of Penicillin or Cephalosporin Hydrolysis. Nat. Commun. 2017, 8 (1), 2242. 10.1038/s41467-017-02339-w.

(4) Zhang, H.; Hao, Q. Crystal Structure of NDM-1 Reveals a Common β-Lactam Hydrolysis Mechanism. FASEB J. Off. Publ. Fed. Am. Soc. Exp. Biol. 2011, 25 (8), 2574–2582. 10.1096/fj.11-184036.

(5) Ma, B.; Fang, C.; Lu, L.; Wang, M.; Xue, X.; Zhou, Y.; Li, M.; Hu, Y.; Luo, X.; Hou, Z. The Antimicrobial Peptide Thanatin Disrupts the Bacterial Outer Membrane and Inactivates the NDM-1 Metallo-β-Lactamase. Nat. Commun. 2019, 10 (1), 3517. 10.1038/s41467-019-11503-3.

(6) Moura, E. C. C. M.; Baeta, T.; Romanelli, A.; Laguri, C.; Martorana, A. M.; Erba, E.; Simorre, J.-P.; Sperandeo, P.; Polissi, A. Thanatin Impairs Lipopolysaccharide Transport Complex Assembly by Targeting LptC-LptA Interaction and Decreasing LptA Stability. Front. Microbiol. 2020, 11, 909. 10.3389/fmicb.2020.00909.

(7) WHO publishes list of bacteria for which new antibiotics are urgently needed. https://www.who.int/news/item/27-02-2017-who-publishes-list-of-bacteria-for-which-new-antibiotics-are-urgently-needed (accessed 2026-02-11).

(8) Tacconelli, E.; Carrara, E.; Savoldi, A.; Harbarth, S.; Mendelson, M.; Monnet, D. L.; Pulcini, C.; Kahlmeter, G.; Kluytmans, J.; Carmeli, Y.; Ouellette, M.; Outterson, K.; Patel, J.; Cavaleri, M.; Cox, E. M.; Houchens, C. R.; Grayson, M. L.; Hansen, P.; Singh, N.; Theuretzbacher, U.; Magrini, N.; Aboderin, A. O.; Al-Abri, S. S.; Jalil, N. A.; Benzonana, N.; Bhattacharya, S.; Brink, A. J.; Burkert, F. R.; Cars, O.; Cornaglia, G.; Dyar, O. J.; Friedrich, A. W.; Gales, A. C.; Gandra, S.; Giske, C. G.; Goff, D. A.; Goossens, H.; Gottlieb, T.; Blanco, M. G.; Hryniewicz, W.; Kattula, D.; Jinks, T.; Kanj, S. S.; Kerr, L.; Kieny, M.-P.; Kim, Y. S.; Kozlov, R. S.; Labarca, J.; Laxminarayan, R.; Leder, K.; Leibovici, L.; Levy-Hara, G.; Littman, J.; Malhotra-Kumar, S.; Manchanda, V.; Moja, L.; Ndoye, B.; Pan, A.; Paterson, D. L.; Paul, M.; Qiu, H.; Ramon-Pardo, P.; Rodríguez-Baño, J.; Sanguinetti, M.; Sengupta, S.; Sharland, M.; Si-Mehand, M.; Silver, L. L.; Song, W.; Steinbakk, M.; Thomsen, J.; Thwaites, G. E.; Meer, J. W. van der; Kinh, N. V.; Vega, S.; Villegas, M. V.; Wechsler-Fördös, A.; Wertheim, H. F. L.; Wesangula, E.; Woodford, N.; Yilmaz, F. O.; Zorzet, A. Discovery, Research, and Development of New Antibiotics: The WHO Priority List of Antibiotic-Resistant Bacteria and Tuberculosis. Lancet Infect. Dis. 2018, 18 (3), 318–327. 10.1016/S1473-3099(17)30753-3.

(9) Vetterli, S. U.; Zerbe, K.; Müller, M.; Urfer, M.; Mondal, M.; Wang, S.-Y.; Moehle, K.; Zerbe, O.; Vitale, A.; Pessi, G.; Eberl, L.; Wollscheid, B.; Robinson, J. A. Thanatin Targets the Intermembrane Protein Complex Required for Lipopolysaccharide Transport in Escherichia Coli. Sci. Adv. 2018, 4 (11), eaau2634. 10.1126/sciadv.aau2634.

(10) Dash, R.; Bhattacharjya, S. Thanatin: An Emerging Host Defense Antimicrobial Peptide with Multiple Modes of Action. Int. J. Mol. Sci. 2021, 22 (4), 1522. 10.3390/ijms22041522.

(11) King, D.; Strynadka, N. Crystal Structure of New Delhi Metallo-β-Lactamase Reveals Molecular Basis for Antibiotic Resistance. Protein Sci. Publ. Protein Soc. 2011, 20 (9), 1484–1491. 10.1002/pro.697.

(12) Honorato, R. V.; Trellet, M. E.; Jiménez-García, B.; Schaarschmidt, J. J.; Giulini, M.; Reys, V.; Koukos, P. I.; Rodrigues, J. P. G. L. M.; Karaca, E.; van Zundert, G. C. P.; Roel-Touris, J.; van Noort, C. W.; Jandová, Z.; Melquiond, A. S. J.; Bonvin, A. M. J. J. The HADDOCK2.4 Web Server for Integrative Modeling of Biomolecular Complexes. Nat. Protoc. 2024, 19 (11), 3219–3241. 10.1038/s41596-024-01011-0.

(13) Dominguez, C.; Boelens, R.; Bonvin, A. M. J. J. HADDOCK: A Protein−Protein Docking Approach Based on Biochemical or Biophysical Information. J. Am. Chem. Soc. 2003, 125 (7), 1731–1737. 10.1021/ja026939x.

(14) Lindorff-Larsen, K.; Piana, S.; Palmo, K.; Maragakis, P.; Klepeis, J. L.; Dror, R. O.; Shaw, D. E. Improved Side-Chain Torsion Potentials for the Amber ff99SB Protein Force Field. Proteins 2010, 78 (8), 1950–1958. 10.1002/prot.22711.

(15) Hornak, V.; Abel, R.; Okur, A.; Strockbine, B.; Roitberg, A.; Simmerling, C. Comparison of Multiple Amber Force Fields and Development of Improved Protein Backbone Parameters. Proteins 2006, 65 (3), 712–725. 10.1002/prot.21123.

(16) Kim, Y.; Cunningham, M. A.; Mire, J.; Tesar, C.; Sacchettini, J.; Joachimiak, A. NDM-1, the Ultimate Promiscuous Enzyme: Substrate Recognition and Catalytic Mechanism. FASEB J. 2013, 27 (5), 1917–1927. 10.1096/fj.12-224014.

(17) Stewart, A. C.; Bethel, C. R.; VanPelt, J.; Bergstrom, A.; Cheng, Z.; Miller, C. G.; Williams, C.; Poth, R.; Morris, M.; Lahey, O.; Nix, J. C.; Tierney, D. L.; Page, R. C.; Crowder, M. W.; Bonomo, R. A.; Fast, W. Clinical Variants of New Delhi Metallo-β-Lactamase Are Evolving to Overcome Zinc Scarcity. ACS Infect. Dis. 2017, 3 (12), 927. 10.1021/acsinfecdis.7b00128.

(18) Sivarajan, V.; Ganesh, A. V.; Subramani, P.; Ganesapandi, P.; Sivanandan, R. N.; Prakash, S.; Manikandan, N.; Dharmarajan, A.; Arfuso, F.; Warrier, S.; Raj, M.; Perumal, K. Prevalence and Genomic Insights of Carbapenem Resistant and ESBL Producing Multidrug Resistant Escherichia Coli in Urinary Tract Infections. Sci. Rep. 2025, 15 (1), 2541. 10.1038/s41598-024-84754-w.

(19) Johnson, A. P.; Woodford, N. Global Spread of Antibiotic Resistance: The Example of New Delhi Metallo-β-Lactamase (NDM)-Mediated Carbapenem Resistance. J. Med. Microbiol. 2013, 62 (Pt 4), 499–513. 10.1099/jmm.0.052555-0.

(20) Xia, X.; Song, S.; Zhang, S.; Wang, W.; Zhou, J.; Fan, B.; Li, L.; Dong, H.; Luo, C.; Li, B.; Zhang, X. The Synergy of Thanatin and Cathelicidin-BF-15a3 Combats Escherichia Coli O157:H7. Int. J. Food Microbiol. 2023, 386, 110018. 10.1016/j.ijfoodmicro.2022.110018.

(21) Shepperson, O. A.; Harris, P. W. R.; Brimble, M. A.; Cameron, A. J. Thanatin and Vinyl Sulfide Analogues as Narrow Spectrum Antimicrobial Peptides That Synergise with Polymyxin B. Front. Pharmacol. 2024, 15. 10.3389/fphar.2024.1487338.

(22) Lob, S. H.; Karlowsky, J. A.; Young, K.; Motyl, M. R.; Hawser, S.; Kothari, N. D.; Sahm, D. F. In Vitro Activity of Imipenem-Relebactam against Resistant Phenotypes of Enterobacteriaceae and Pseudomonas Aeruginosa Isolated from Intraabdominal and Urinary Tract Infection Samples – SMART Surveillance Europe 2015–2017. J. Med. Microbiol. 2020, 69 (2), 207–217. 10.1099/jmm.0.001142.

(23) Mansour, H.; Ouweini, A. E. L.; Chahine, E. B.; Karaoui, L. R. Imipenem/Cilastatin/Relebactam: A New Carbapenem β-Lactamase Inhibitor Combination. Am. J. Health-Syst. Pharm. AJHP Off. J. Am. Soc. Health-Syst. Pharm. 2021, 78 (8), 674–683. 10.1093/ajhp/zxab012.

(24) Johnston, B. D.; Thuras, P.; Porter, S. B.; Anacker, M.; VonBank, B.; Vagnone, P. S.; Witwer, M.; Castanheira, M.; Johnson, J. R. Activity of Imipenem-Relebactam against Carbapenem-Resistant Escherichia Coli Isolates from the United States in Relation to Clonal Background, Resistance Genes, Coresistance, and Region. Antimicrob. Agents Chemother. 2020, 64 (5), e02408–19. 10.1128/AAC.02408-19.

