## Supplementary Information for "Allosteric Inhibition of NDM-1 by Thanatin Preserves the Di-Zinc Center While Restricting Dynamics"

➢ Supplementary Figures

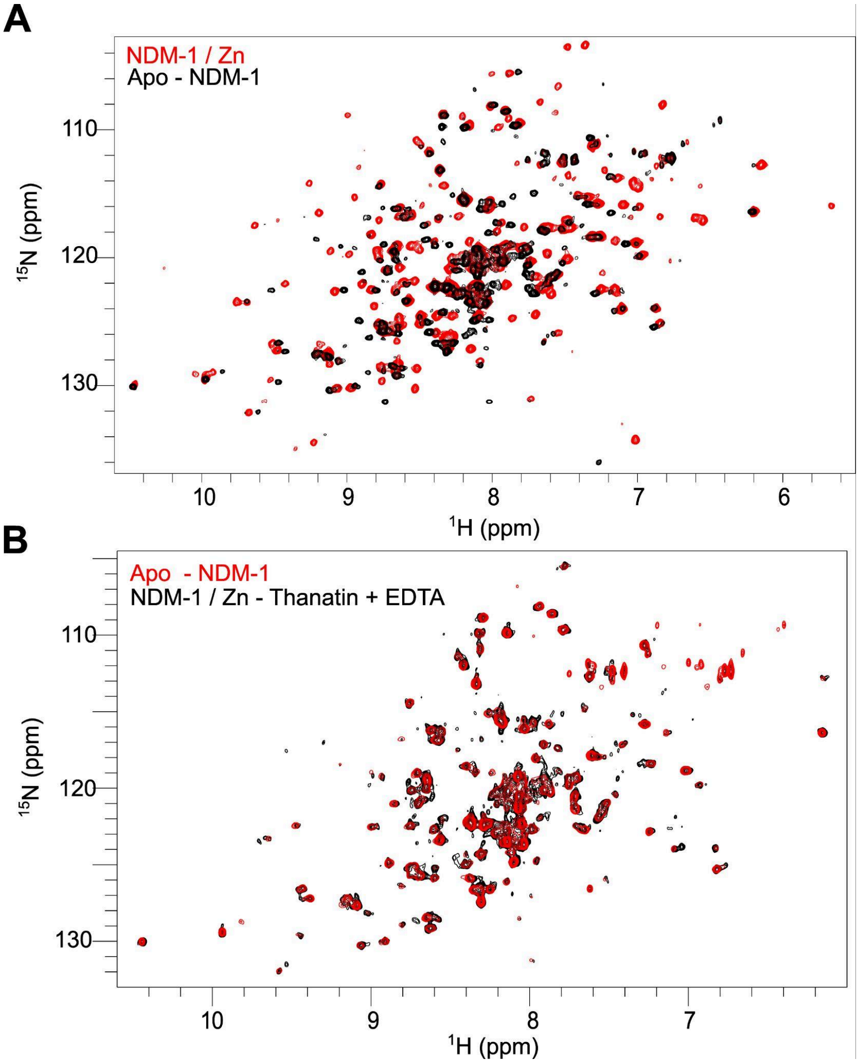

**Figure SI1: EDTA-induced zinc dissociation from the NDM-1–thanatin complex restores the apo-NDM-1 HSQC signature.**

(A) Overlay of $^{1}$H-$^{15}$N HSQC spectra of apo-NDM-1 (black) and $Zn^{2+}$-bound NDM-1 (red), showing widespread chemical shift perturbations upon zinc binding consistent with global conformational stabilization of the enzyme. (B) Overlay of $^{1}$H-$^{15}$N HSQC spectra of the NDM-1 (4 eq $Zn^{2+}$)–thanatin complex after EDTA addition (black) and apo-NDM-1 (red). Removal of zinc from the ternary complex restores an apo-like spectral pattern, indicating complete zinc dissociation.

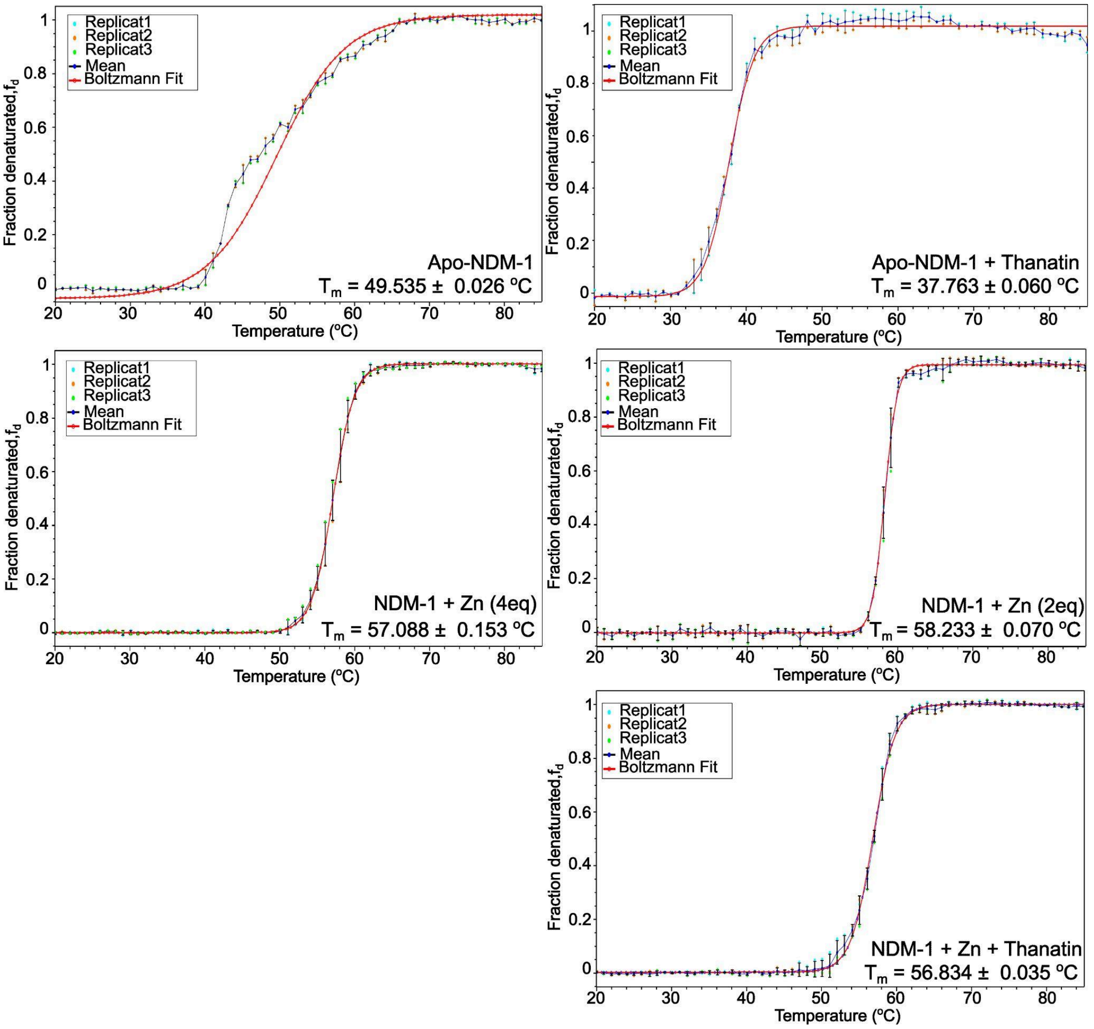

**<u>Figure SI2: Thermal denaturation of NDM-1 monitored by CD spectroscopy.</u>**

(A) Apo-NDM-1 (Tm = 49.5 ± 0.03 °C, n = 3). (B) Thanatin-bound apo-NDM-1 (Tm = 37.8 ± 0.06 °C, n = 3). (C) $Zn^{2+}$-bound NDM-1 at 2 eq and 4 eq (Tm = 58.2 ± 0.1 °C and 57.1 ± 0.2 °C, n = 3). (D) Thanatin-bound Zn-NDM-1 (Tm = 57.5 ± 1.1 °C, n = 3). Curves represent mean values (n = 3); shaded regions indicate ± SD. Fits were obtained using a Boltzmann sigmoidal model.

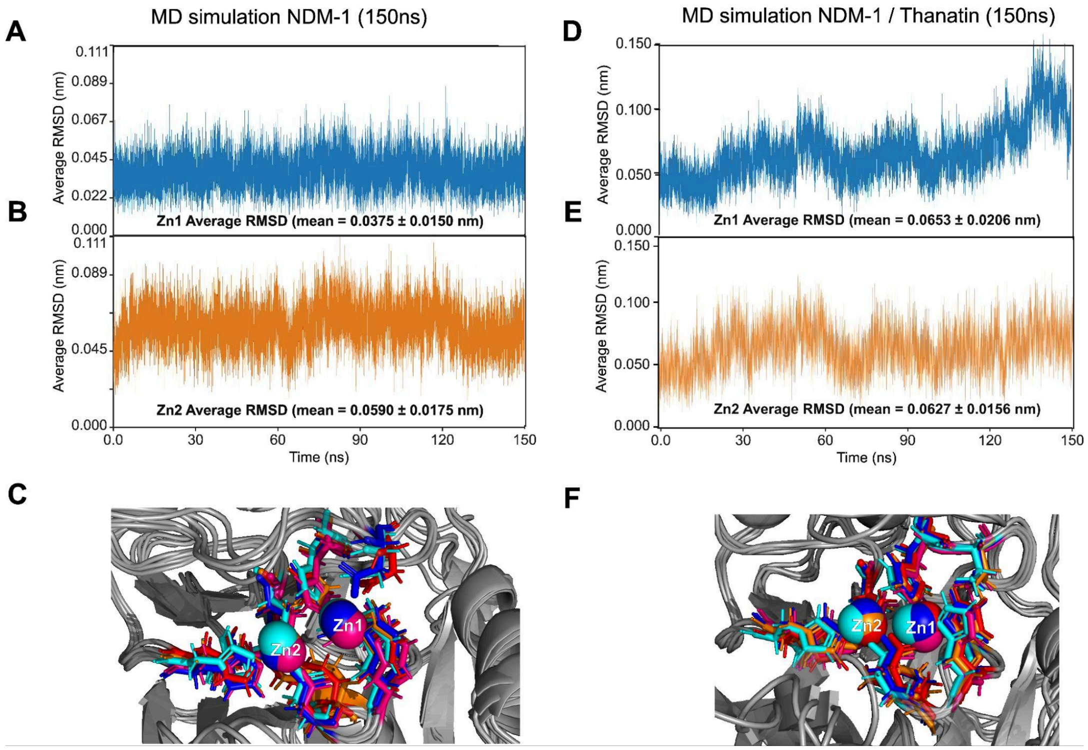

**Figure SI3: Stability of the NDM-1 Catalytic Di-zinc Center throughout Molecular Dynamics Simulations.**

Top and middle panels show the Root-Mean-Square Deviation (RMSD) of the catalytic zinc ions over 150 ns of production MD. (A–B) Stability of Zn1 and Zn2 in the native holo-NDM-1 (PDB: 3ZR9) control, showing mean RMSD values of 0.0375 ± 0.0150 nm and 0.0590 ± 0.0175 nm, respectively. (D–E) Stability of the zinc ions in the NDM-1/Thanatin complex. The preservation of low RMSD values (< 1 Å) in the presence of the peptide demonstrates that Thanatin binding does not perturb the metal coordination sphere. (C & F) Structural overlays of the NDM-1 active site (Zn1 in blue; Zn2 in pink) at t = 0 ns (blue), t = 20 ns (red), t = 80 ns (cyan), and t = 150 ns (orange) illustrating the rigorous maintenance of the coordination geometry between the zinc ions and their respective protein ligands (His120, His122, His189 for Zn1; Asp124, Cys208, His250 for Zn2). Collectively, these simulations support the zinc-retaining allosteric mechanism observed by NMR and preclude a metal-displacement model for Thanatin-mediated inhibition.

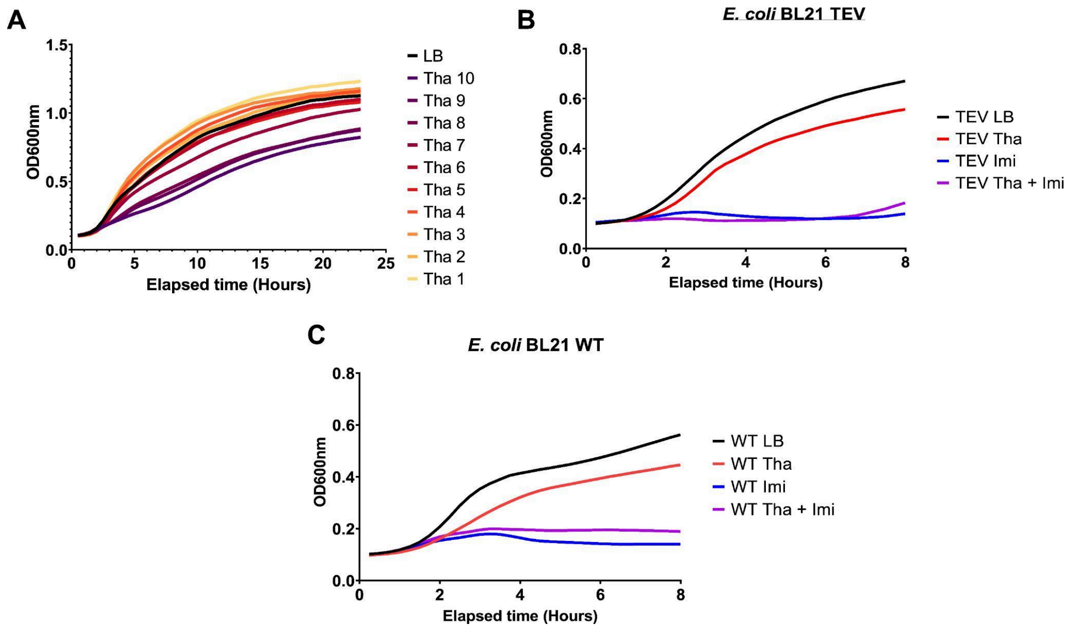

**Figure SI4: Control Growth Dynamics and Synergistic Potential of Thanatin with Imipenem**

(A) Representative growth curves ($OD_{600}$) of *E. coli* BL21 (DE3) expressing NDM-1 in the presence of varying concentrations of thanatin (1 - 10 µM). The dose-response profile over 24 h indicates that thanatin lacks potent primary antimicrobial activity against this resistant strain at the levels tested. (B) and (C) Comparative growth kinetics of *E. coli* BL21 (DE3) harboring the NDM-1 expression plasmid and wild-type (WT) control. Cultures were induced with 1 mM IPTG and monitored over 8 h at 37 °C. Treatment groups include: LB control (black), 6 µM thanatin (red), 1 µg/mL imipenem (blue), and the combination of 6 µM thanatin and 1 µg/mL imipenem (purple). These data confirm that 6 µM thanatin does not significantly inhibit bacterial proliferation as a standalone agent, supporting its role as a non-toxic allosteric modulator that potentiates imipenem activity. The dramatic growth suppression observed only in the combination group (purple) demonstrates the successful reversal of NDM-1 mediated carbapenem resistance.

➢ Supplementary Tables

**Table SI1: List of intermolecular NOE between NDM-1 residues and Thanatin's protons.**

| NDM-1 residues | Proton residues from Thanatin |
|---|---|
| Gln13 | 4.292 ; 0.802 ppm |
| Met14 | 4.320 ppm |
| Thr16 | 4.410 ppm |
| Asp18 | 4.361 ; 4.818 ppm |
| Arg20 | 1.519 ppm |
| Phe21 | 4.840 ; 0.733 ppm |
| Leu24 | 4.966 ; 1.852 ppm |
| Val25 | 4.966 ; 1.852 ppm |
| Thr37 | 5.430 ppm |
| Ser38 | 5.430 ppm |
| Asp41 | 3.99 ppm |
| Met42 | 4.12 ppm |
| Phe45 | 8.108 ; 3.032 ; 2.645 ppm |
| Gly46 | 8.135 ; 2.706 ; 2.988 ppm |
| Ala67 | 5.598 ; 4.624 ppm |
| Trp68 | 5.492 ppm |
| Asp71 | 5.651 ppm |
| Ile76 | 2.127 ppm |
| Trp79 | 5.747 ppm |
| Glu83 | 5.311 ppm |
| Asp99 | 6.992 ppm |
| Gly 103 | 5.591 ppm |
| Ala116 | 5.167 ppm |

| | |
|---|---|
| Gln122 | 4.100 ppm |
| Leu123 | 4.091 ppm |
| Gln126 | 4.643 ; 2.351 ppm |
| Thr165 | 3.826 ; 0.375 ppm |

**Table SI2: Thermal denaturation parameters for NDM-1 under apo-NDM-1, apo-NDM-1 + thanatin, $Zn^{2+}$-bound (2 eq and 4 eq), and $Zn^{2+}$ + thanatin conditions.**

Apparent melting temperatures ($T_m$) were extracted from Boltzmann sigmoidal fits of normalized CD data at 222 nm. Standard deviations reflect replicate variability (n = 3); $R^2$ values indicate goodness of fit. Apo-NDM-1 unfolds broadly and non-cooperatively ($T_m$ = 49.5 ± 0.03 °C; slope = 0.058 ± 0.001), while $Zn^{2+}$ binding increases thermal stability and sharpens the transition ($T_m$ = 58.2 ± 0.07 °C at 2 eq $Zn^{2+}$; 57.1 ± 0.15 °C at 4 eq $Zn^{2+}$). Thanatin binding to apo-NDM-1 lowers $T_m$ (37.8 ± 0.06 °C) while increasing cooperativity (slope = 0.158 ± 0.005), consistent with Zn-independent association. In $Zn^{2+}$-bound NDM-1, thanatin does not alter $T_m$ (56.8 ± 0.04 °C vs 57.1 ± 0.15 °C), indicating $Zn^{2+}$ retention and preserved global fold. Full replicate traces and fit parameters are shown in Figure SI2.

| | $T_m$ (ºC) | $R^2$ | Slope |
|---|---|---|---|
| NDM-1 alone | 49.535 ± 0.026 | 0.9986 | 0.058 ± 0.001 |
| NDM-1 + Thanatin | 37.763 ± 0.060 | 0.9998 | 0.158 ± 0.005 |
| NDM-1 $Zn^{2+}$(2eq) | 58.223 ± 0.070 | 0.9997 | 0.311 ± 0.014 |
| NDM-1 $Zn^{2+}$(4eq) | 57.088 ± 0.153 | 0.9999 | 0.173 ± 0.007 |
| NDM-1/$Zn^{2+}$/Thanatin | 56.834 ± 0.035 | 0.9997 | 0.169 ± 0.004 |

**Table SI3: HADDOCK Score and Intermolecular Energy Statistics for Docking Clusters.**

| Cluster | haddock-score | sd | rmsd | sd | rmsd-Emin | sd | Nstruc | Einter | sd |
|---|---|---|---|---|---|---|---|---|---|
| file.nam_clust4 | -95.52 | 8.185 | 0.994 | 0.59 | 0.994 | 0.59 | 11 | -417.06 | 60.58 |
| file.nam_clust7 | -84.527 | 10.286 | 0.903 | 0.535 | 3.619 | 0.314 | 6 | -220.33 | 40.11 |
| file.nam_clust6 | -81.065 | 7 | 0.627 | 0.376 | 5.234 | 0.207 | 7 | -170.22 | 49.81 |
| file.nam_clust2 | -78.112 | 3.676 | 1.222 | 0.707 | 3.031 | 0.269 | 21 | -277.21 | 16.68 |
| file.nam_clust3 | -74.568 | 6.45 | 0.965 | 0.719 | 2.754 | 0.217 | 15 | -167.35 | 56.39 |
| file.nam_clust1 | -74.524 | 2.87 | 0.76 | 0.501 | 4.4 | 0.05 | 29 | -271.75 | 61.29 |
| file.nam_clust12 | -72.505 | 4.614 | 0.848 | 0.589 | 4.565 | 0.064 | 4 | -203.06 | 47.05 |
| file.nam_clust5 | -72.405 | 4.964 | 1.37 | 0.85 | 4.676 | 0.159 | 9 | -129.29 | 39.34 |
| file.nam_clust10 | -69.059 | 10.28 | 1.175 | 0.71 | 4.672 | 0.049 | 5 | -196.72 | 91.38 |
| file.nam_clust9 | -68.961 | 8.411 | 0.565 | 0.374 | 2.655 | 0.048 | 5 | -221.77 | 38.94 |
| file.nam_clust11 | -66.62 | 7.53 | 0.706 | 0.45 | 3.521 | 0.235 | 5 | -160.13 | 48.02 |
| file.nam_clust8 | -65.907 | 1.679 | 0.685 | 0.513 | 4.401 | 0.155 | 6 | -135.82 | 37.82 |
| file.nam_clust13 | -60.915 | 6.305 | 0.858 | 0.498 | 3.973 | 0.091 | 4 | -49.94 | 70.22 |
| file.nam_clust14 | -60.833 | 9.427 | 1.109 | 0.773 | 4.901 | 0.226 | 4 | -165.21 | 45.66 |

**Table SI4: List of atom-atom interactions across NDM-1/thanatin interface calculated from PDBsum generator**

List of atom-atom interactions across protein-protein interface

Hydrogen bonds

|  | <----- A T O M 1 -----> |  |  |  |  | <----- A T O M 2 -----> |  |  |  |  |  |
|---|---|---|---|---|---|---|---|---|---|---|---|
| 1. | 254 | OD1 | ASP | 66 | A | <--> | 2216 | NE | ARG | 14 | B | 2.66 |
| 2. | 255 | OD2 | ASP | 66 | A | <--> | 2222 | NH2 | ARG | 14 | B | 2.60 |
| 3. | 515 | OD1 | ASP | 96 | A | <--> | 2248 | NZ | LYS | 17 | B | 2.57 |
| 4. | 754 | O | ALA | 121 | A | <--> | 2115 | NZ | LYS | 4 | B | 2.72 |
| 5. | 835 | OD2 | ASP | 130 | A | <--> | 2282 | NH1 | ARG | 20 | B | 2.59 |
| 6. | 1026 | O | GLU | 152 | A | <--> | 2115 | NZ | LYS | 4 | B | 2.77 |
| 7. | 1023 | OE1 | GLU | 152 | A | <--> | 2115 | NZ | LYS | 4 | B | 2.55 |
| 8. | 1646 | OD2 | ASP | 223 | A | <--> | 2102 | NZ | LYS | 3 | B | 2.59 |

Non-bonded contacts

|  | <----- A T O M 1 -----> |  |  |  |  | <----- A T O M 2 -----> |  |  |  |  |  |
|---|---|---|---|---|---|---|---|---|---|---|---|
| 1. | 64 | CB | ASP | 48 | A | <--> | 2225 | C | ARG | 14 | B | 3.76 |
| 2. | 64 | CB | ASP | 48 | A | <--> | 2226 | O | ARG | 14 | B | 3.38 |
| 3. | 65 | CG | ASP | 48 | A | <--> | 2225 | C | ARG | 14 | B | 3.82 |
| 4. | 65 | CG | ASP | 48 | A | <--> | 2226 | O | ARG | 14 | B | 3.88 |
| 5. | 65 | CG | ASP | 48 | A | <--> | 2213 | CB | ARG | 14 | B | 3.73 |
| 6. | 66 | OD1 | ASP | 48 | A | <--> | 2230 | CB | THR | 15 | B | 3.68 |

| | | | | | | | | | | | | |
|---|---|---|---|---|---|---|---|---|---|---|---|---|
| 7. | 67 | OD2 | ASP | 48 | A | <--> | 2213 | CB | ARG | 14 | B | 3.08 |
| 8. | 239 | O | TYR | 64 | A | <--> | 2214 | CG | ARG | 14 | B | 3.50 |
| 9. | 253 | CG | ASP | 66 | A | <--> | 2216 | NE | ARG | 14 | B | 3.50 |
| 10. | 253 | CG | ASP | 66 | A | <--> | 2218 | CZ | ARG | 14 | B | 3.87 |
| 11. | 253 | CG | ASP | 66 | A | <--> | 2222 | NH2 | ARG | 14 | B | 3.32 |
| 12. | 254 | OD1 | ASP | 66 | A | <--> | 2214 | CG | ARG | 14 | B | 3.59 |
| 13. | 254 | OD1 | ASP | 66 | A | <--> | 2215 | CD | ARG | 14 | B | 3.67 |
| 14. | 254 | OD1 | ASP | 66 | A | <--> | 2216 | NE | ARG | 14 | B | 2.66 |
| 15. | 254 | OD1 | ASP | 66 | A | <--> | 2218 | CZ | ARG | 14 | B | 3.39 |
| 16. | 254 | OD1 | ASP | 66 | A | <--> | 2222 | NH2 | ARG | 14 | B | 3.29 |
| 17. | 255 | OD2 | ASP | 66 | A | <--> | 2216 | NE | ARG | 14 | B | 3.60 |
| 18. | 255 | OD2 | ASP | 66 | A | <--> | 2218 | CZ | ARG | 14 | B | 3.52 |
| 19. | 255 | OD2 | ASP | 66 | A | <--> | 2222 | NH2 | ARG | 14 | B | 2.60 |
| 20. | 495 | CB | THR | 94 | A | <--> | 2265 | CG | GLN | 19 | B | 3.46 |
| 21. | 495 | CB | THR | 94 | A | <--> | 2266 | CD | GLN | 19 | B | 3.46 |
| 22. | 495 | CB | THR | 94 | A | <--> | 2267 | OE1 | GLN | 19 | B | 3.27 |
| 23. | 496 | OG1 | THR | 94 | A | <--> | 2188 | ND2 | ASN | 12 | B | 3.63 |
| 24. | 496 | OG1 | THR | 94 | A | <--> | 2266 | CD | GLN | 19 | B | 3.81 |
| 25. | 496 | OG1 | THR | 94 | A | <--> | 2267 | OE1 | GLN | 19 | B | 3.15 |
| 26. | 498 | CG2 | THR | 94 | A | <--> | 2169 | CE2 | TYR | 10 | B | 3.69 |
| 27. | 498 | CG2 | THR | 94 | A | <--> | 2171 | OH | TYR | 10 | B | 3.41 |
| 28. | 498 | CG2 | THR | 94 | A | <--> | 2266 | CD | GLN | 19 | B | 3.82 |
| 29. | 498 | CG2 | THR | 94 | A | <--> | 2267 | OE1 | GLN | 19 | B | 3.80 |
| 30. | 501 | N | ASP | 95 | A | <--> | 2265 | CG | GLN | 19 | B | 3.57 |
| 31. | 501 | N | ASP | 95 | A | <--> | 2289 | O | ARG | 20 | B | 3.79 |
| 32. | 504 | CB | ASP | 95 | A | <--> | 2264 | CB | GLN | 19 | B | 3.77 |
| 33. | 504 | CB | ASP | 95 | A | <--> | 2289 | O | ARG | 20 | B | 3.22 |
| 34. | 510 | N | ASP | 96 | A | <--> | 2264 | CB | GLN | 19 | B | 3.58 |
| 35. | 510 | N | ASP | 96 | A | <--> | 2265 | CG | GLN | 19 | B | 3.68 |
| 36. | 513 | CB | ASP | 96 | A | <--> | 2233 | CG2 | THR | 15 | B | 3.76 |
| 37. | 513 | CB | ASP | 96 | A | <--> | 2267 | OE1 | GLN | 19 | B | 3.22 |
| 38. | 514 | CG | ASP | 96 | A | <--> | 2246 | CD | LYS | 17 | B | 3.75 |
| 39. | 514 | CG | ASP | 96 | A | <--> | 2248 | NZ | LYS | 17 | B | 3.03 |
| 40. | 514 | CG | ASP | 96 | A | <--> | 2258 | O | CYS | 18 | B | 3.84 |
| 41. | 514 | CG | ASP | 96 | A | <--> | 2264 | CB | GLN | 19 | B | 3.69 |
| 42. | 514 | CG | ASP | 96 | A | <--> | 2267 | OE1 | GLN | 19 | B | 3.83 |
| 43. | 515 | OD1 | ASP | 96 | A | <--> | 2246 | CD | LYS | 17 | B | 3.36 |

| | | | | | | | | | | | | |
|---|---|---|---|---|---|---|---|---|---|---|---|---|
| 44. | 515 | OD1 | ASP | 96 | A | <--> | 2247 | CE | LYS | 17 | B | 3.30 |
| 45. | 515 | OD1 | ASP | 96 | A | <--> | 2248 | NZ | LYS | 17 | B | 2.57 |
| 46. | 516 | OD2 | ASP | 96 | A | <--> | 2252 | C | LYS | 17 | B | 3.63 |
| 47. | 516 | OD2 | ASP | 96 | A | <--> | 2253 | O | LYS | 17 | B | 3.85 |
| 48. | 516 | OD2 | ASP | 96 | A | <--> | 2244 | CB | LYS | 17 | B | 3.47 |
| 49. | 516 | OD2 | ASP | 96 | A | <--> | 2245 | CG | LYS | 17 | B | 3.53 |
| 50. | 516 | OD2 | ASP | 96 | A | <--> | 2246 | CD | LYS | 17 | B | 3.58 |
| 51. | 516 | OD2 | ASP | 96 | A | <--> | 2247 | CE | LYS | 17 | B | 3.74 |
| 52. | 516 | OD2 | ASP | 96 | A | <--> | 2248 | NZ | LYS | 17 | B | 2.76 |
| 53. | 516 | OD2 | ASP | 96 | A | <--> | 2254 | N | CYS | 18 | B | 3.66 |
| 54. | 516 | OD2 | ASP | 96 | A | <--> | 2257 | C | CYS | 18 | B | 3.17 |
| 55. | 516 | OD2 | ASP | 96 | A | <--> | 2258 | O | CYS | 18 | B | 2.77 |
| 56. | 516 | OD2 | ASP | 96 | A | <--> | 2261 | N | GLN | 19 | B | 3.67 |
| 57. | 516 | OD2 | ASP | 96 | A | <--> | 2263 | CA | GLN | 19 | B | 3.85 |
| 58. | 516 | OD2 | ASP | 96 | A | <--> | 2264 | CB | GLN | 19 | B | 3.21 |
| 59. | 516 | OD2 | ASP | 96 | A | <--> | 2266 | CD | GLN | 19 | B | 3.70 |
| 60. | 516 | OD2 | ASP | 96 | A | <--> | 2267 | OE1 | GLN | 19 | B | 3.47 |
| 61. | 524 | CD | GLN | 97 | A | <--> | 2188 | ND2 | ASN | 12 | B | 3.60 |
| 62. | 526 | NE2 | GLN | 97 | A | <--> | 2188 | ND2 | ASN | 12 | B | 3.47 |
| 63. | 526 | NE2 | GLN | 97 | A | <--> | 2213 | CB | ARG | 14 | B | 3.84 |
| 64. | 753 | C | ALA | 121 | A | <--> | 2115 | NZ | LYS | 4 | B | 3.79 |
| 65. | 754 | O | ALA | 121 | A | <--> | 2114 | CE | LYS | 4 | B | 3.53 |
| 66. | 754 | O | ALA | 121 | A | <--> | 2115 | NZ | LYS | 4 | B | 2.72 |
| 67. | 757 | CA | HIS | 122 | A | <--> | 2114 | CE | LYS | 4 | B | 3.56 |
| 68. | 757 | CA | HIS | 122 | A | <--> | 2115 | NZ | LYS | 4 | B | 3.85 |
| 69. | 765 | C | HIS | 122 | A | <--> | 2114 | CE | LYS | 4 | B | 3.88 |
| 70. | 761 | CD2 | HIS | 122 | A | <--> | 2114 | CE | LYS | 4 | B | 3.77 |
| 71. | 767 | N | GLN | 123 | A | <--> | 2113 | CD | LYS | 4 | B | 3.48 |
| 72. | 767 | N | GLN | 123 | A | <--> | 2114 | CE | LYS | 4 | B | 3.44 |
| 73. | 770 | CB | GLN | 123 | A | <--> | 2113 | CD | LYS | 4 | B | 3.62 |
| 74. | 774 | NE2 | GLN | 123 | A | <--> | 2171 | OH | TYR | 10 | B | 3.72 |
| 75. | 814 | O | GLY | 127 | A | <--> | 2289 | O | ARG | 20 | B | 3.17 |
| 76. | 814 | O | GLY | 127 | A | <--> | 2292 | CA | MET | 21 | B | 3.68 |
| 77. | 814 | O | GLY | 127 | A | <--> | 2297 | C | MET | 21 | B | 3.71 |
| 78. | 833 | CG | ASP | 130 | A | <--> | 2278 | CD | ARG | 20 | B | 3.86 |
| 79. | 833 | CG | ASP | 130 | A | <--> | 2282 | NH1 | ARG | 20 | B | 3.53 |
| 80. | 834 | OD1 | ASP | 130 | A | <--> | 2282 | NH1 | ARG | 20 | B | 3.84 |

| | | | | | | | | | | | | |
|---|---|---|---|---|---|---|---|---|---|---|---|---|
| 81. | 835 | OD2 | ASP | 130 | A | <--> | 2277 | CG | ARG | 20 | B | 3.89 |
| 82. | 835 | OD2 | ASP | 130 | A | <--> | 2278 | CD | ARG | 20 | B | 2.95 |
| 83. | 835 | OD2 | ASP | 130 | A | <--> | 2279 | NE | ARG | 20 | B | 3.66 |
| 84. | 835 | OD2 | ASP | 130 | A | <--> | 2281 | CZ | ARG | 20 | B | 3.54 |
| 85. | 835 | OD2 | ASP | 130 | A | <--> | 2282 | NH1 | ARG | 20 | B | 2.59 |
| 86. | 1019 | CA | GLU | 152 | A | <--> | 2115 | NZ | LYS | 4 | B | 3.79 |
| 87. | 1025 | C | GLU | 152 | A | <--> | 2115 | NZ | LYS | 4 | B | 3.65 |
| 88. | 1026 | O | GLU | 152 | A | <--> | 2112 | CG | LYS | 4 | B | 3.13 |
| 89. | 1026 | O | GLU | 152 | A | <--> | 2113 | CD | LYS | 4 | B | 3.69 |
| 90. | 1026 | O | GLU | 152 | A | <--> | 2114 | CE | LYS | 4 | B | 3.78 |
| 91. | 1026 | O | GLU | 152 | A | <--> | 2115 | NZ | LYS | 4 | B | 2.77 |
| 92. | 1020 | CB | GLU | 152 | A | <--> | 2115 | NZ | LYS | 4 | B | 3.08 |
| 93. | 1021 | CG | GLU | 152 | A | <--> | 2115 | NZ | LYS | 4 | B | 3.62 |
| 94. | 1022 | CD | GLU | 152 | A | <--> | 2101 | CE | LYS | 3 | B | 3.50 |
| 95. | 1022 | CD | GLU | 152 | A | <--> | 2115 | NZ | LYS | 4 | B | 3.38 |
| 96. | 1023 | OE1 | GLU | 152 | A | <--> | 2101 | CE | LYS | 3 | B | 3.10 |
| 97. | 1023 | OE1 | GLU | 152 | A | <--> | 2102 | NZ | LYS | 3 | B | 3.60 |
| 98. | 1023 | OE1 | GLU | 152 | A | <--> | 2114 | CE | LYS | 4 | B | 3.22 |
| 99. | 1023 | OE1 | GLU | 152 | A | <--> | 2115 | NZ | LYS | 4 | B | 2.55 |
| 100. | 1024 | OE2 | GLU | 152 | A | <--> | 2101 | CE | LYS | 3 | B | 3.11 |
| 101. | 1024 | OE2 | GLU | 152 | A | <--> | 2102 | NZ | LYS | 3 | B | 3.81 |
| 102. | 1031 | O | GLY | 153 | A | <--> | 2294 | CG | MET | 21 | B | 3.56 |
| 103. | 1031 | O | GLY | 153 | A | <--> | 2295 | SD | MET | 21 | B | 3.67 |
| 104. | 1036 | CG | MET | 154 | A | <--> | 2115 | NZ | LYS | 4 | B | 3.77 |
| 105. | 1038 | CE | MET | 154 | A | <--> | 2111 | CB | LYS | 4 | B | 3.83 |
| 106. | 1038 | CE | MET | 154 | A | <--> | 2113 | CD | LYS | 4 | B | 3.53 |
| 107. | 1038 | CE | MET | 154 | A | <--> | 2293 | CB | MET | 21 | B | 3.61 |
| 108. | 1038 | CE | MET | 154 | A | <--> | 2294 | CG | MET | 21 | B | 3.69 |
| 109. | 1038 | CE | MET | 154 | A | <--> | 2295 | SD | MET | 21 | B | 3.66 |
| 110. | 1046 | CG2 | VAL | 155 | A | <--> | 2298 | O | MET | 21 | B | 3.89 |
| 111. | 1644 | CG | ASP | 223 | A | <--> | 2102 | NZ | LYS | 3 | B | 3.46 |
| 112. | 1646 | OD2 | ASP | 223 | A | <--> | 2101 | CE | LYS | 3 | B | 3.84 |
| 113. | 1646 | OD2 | ASP | 223 | A | <--> | 2102 | NZ | LYS | 3 | B | 2.59 |

Salt bridges

|  | <----- A T O M 1 -----> |  |  |  |  |  | <----- A T O M 2 -----> |  |  |  |  |  |
|---|---|---|---|---|---|---|---|---|---|---|---|---|
| 1. | 254 | OD1 | ASP | 66 | A | <--> | 2222 | NH2 | ARG | 14 | B | 2.60 |

| | | | | | | | | | | | | |
|---|---|---|---|---|---|---|---|---|---|---|---|---|
| 2. | 515 | OD1 | ASP | 96 | A | <--> | 2248 | NZ | LYS | 17 | B | 2.57 |
| 3. | 835 | OD2 | ASP | 130 | A | <--> | 2282 | NH1 | ARG | 20 | B | 2.59 |
| 4. | 1023 | OE1 | GLU | 152 | A | <--> | 2102 | NZ | LYS | 3 | B | 3.60 |
| 5. | 1023 | OE1 | GLU | 152 | A | <--> | 2115 | NZ | LYS | 4 | B | 2.55 |
| 6. | 1645 | OD1 | ASP | 223 | A | <--> | 2102 | NZ | LYS | 3 | B | 2.59 |

Number of salt bridges: 6

Number of hydrogen bonds: 8

Number of non-bonded contacts: 113

➢ Experimental Methods

**Expression and purification of NDM-1 metallo-β-lactamases (MBLs).**

The gene encoding NDM-1 (amino acids G29–R270; Uniprot ID: C7C422) was subcloned into the pET28a(+) TEV expression vector, including an N-terminal $His_6$ tag for purification and a TEV protease cleavage site for tag removal.

$^{15}N$- and $^{13}C/^{15}N$-labeled NDM-1 were overexpressed in *E. coli* BL21 cells. A 1 L LB culture containing 40 μg/mL kanamycin was inoculated with 10 mL of a fresh overnight preculture and grown at 37 °C to $OD_{600}$ ≈ 1.0. Cells were harvested (5000 rpm, 10 min), resuspended in 1 L pre-warmed M9 minimal medium (6 g/L $Na_2HPO_4$, 3 g/L $KH_2PO_4$, 0.5 g/L NaCl, 1 mM $MgSO_4$, 0.1 mM $CaCl_2$) with 40 μg/mL kanamycin and either (a) 2 g/L [$^{13}C$]glucose and 1 g/L [$^{15}N$]$NH_4Cl$ for double labeling or (b) 2 g/L glucose and 1 g/L [$^{15}N$]$NH_4Cl$ for single labeling. Cultures were grown at 37 °C to $OD_{600}$ ≈ 0.6, then induced with 1 mM IPTG and incubated at 25 °C for 20 h. Cells were harvested (5000 rpm, 30 min), resuspended in 30 mL buffer A (50 mM sodium phosphate, 0.2 M NaCl, pH 7.5) containing 0.5 mg/mL lysozyme, 100 μg/mL DNase, and protease inhibitor cocktail (Roche), and lysed by sonication (15 × 10 s pulses at 4 °C). Cell debris was removed by centrifugation (10,000 rpm, 1 h). The supernatant was applied to a 5 mL HisTrap HP column (Cytiva) equilibrated in buffer A with 25 mM imidazole. After washing with 20 mL of the same buffer, protein was eluted with buffer A containing 0.5 M imidazole.

The eluted fraction was dialyzed overnight at 4 °C against 0.1 M Tris-HCl, pH 8.0, and digested with TEV protease (1:100 protein:TEV) in the same buffer supplemented with 0.5 mM EDTA and 1 mM DTT at 4 °C overnight. A second HisTrap HP column was used to separate NDM-1 from the cleaved $His_6$ tag: NDM-1 appeared in the flow-through and wash fractions (buffer A with 25 mM imidazole), while the tag was eluted with 0.5 M imidazole. The purified

NDM-1 fraction was concentrated to 500 µM using a 10 kDa centrifugal concentrator and dialyzed against 0.1 mM Bis-Tris-HCl, 150 mM NaCl, pH 7.0. Aliquots were stored at −80 °C until use.

**NMR studies**

All NMR spectra were acquired on a Bruker Avance III 700 MHz spectrometer equipped with a 5 mm z-gradient TCI (H/C/N) cryoprobe at 30 °C. Data were processed with TopSpin 3.5 (Bruker) and analyzed using CcpNmr 2.4.2 (http://www.ccpn.ac.uk) [1]. All NDM-1 spectra were recorded in phosphate-free Bis-Tris-HCl buffer to avoid precipitation of low-solubility zinc phosphate [2].

**NDM-1 NMR Sample Preparation:*

Purified NDM-1 was prepared in buffer B: 10 mM Bis-Tris-HCl, 150 mM NaCl, pH 7.0. To ensure full zinc saturation, $ZnCl_2$ (100 mM stock) was added to a final stoichiometry of ≈4 equivalents per monomer; saturation was confirmed by stabilization of the $^1H$–$^{15}N$ HSQC spectrum. The N-terminal $His_6$ tag was removed prior to NMR acquisition to minimize aggregation.

**$^1H$–$^{15}N$ HSQC Titration Experiments:*

$^1H$–$^{15}N$ TROSY-HSQC spectra were acquired at 150 µM NDM-1 with incremental additions of unlabeled thanatin (NovoPro; supplied without TFA) dissolved in buffer B (10 mM stock). NDM-1 was titrated with 0–10 equivalents of thanatin. Each spectrum was collected with 240 scans per increment (spectral widths: 12.62 and 2.4 kHz for $^1H$ and $^{15}N$, respectively; 128 points in the indirect dimension). Weighted chemical shift changes (Δδ) were calculated as:

$$\Delta\delta = [\,(\Delta\delta H)^2 + 0.14\cdot(\Delta\delta N)^2\,]^{1/2}$$

where ΔδH and ΔδN are the chemical shift changes in the $^1H$ and $^{15}N$ dimensions, respectively

*$^{15}N$ Relaxation Measurements:*

$R_1$ and $R_2$ relaxation rates were measured for di-$Zn^{2+}$-bound NDM-1 and its complex with 10 equivalents of thanatin using TROSY-based pseudo-3D experiments. $R_1$ datasets were collected at relaxation delays of 20 (duplicated), 150, 300, 500, 600, 800, 1000, and 1500 ms. $R_2$ datasets used delays of 16.96 (duplicated), 33.92, 50.88, 67.84, 101.76, 135.68, 169.60, and 203.52 ms. Acquisition times were 104.44 and 25.76 ms in the $^{1}H$ and $^{15}N$ dimensions, respectively, with 48 scans per increment. Relaxation rates were obtained by fitting peak intensities to mono-exponential decays using CCPNMR Analysis.

Backbone resonance assignments for NDM-1 were obtained from the Biological Magnetic Resonance Bank (BMRB: 26950) and confirmed using standard triple-resonance experiments (HNCA, HN(CO)CA, HNCO, HN(CA)CO, HNCACB). Protein concentrations for assignment experiments were optimized for data quality and stability; NDM-1 was prepared at 1 mM.

*$^{15}N$ -$^{13}C$ filtered HSQC NOESY:*

$^{15}N/^{13}C$-filtered NOESY-HSQC spectra were recorded to identify intermolecular NOEs between thanatin and NDM-1 by selectively suppressing $^{1}H$ signals from $^{15}N/^{13}C$-labeled NDM-1. A mixing time of 150 ms was used to optimize the detection of short-range intermolecular NOEs. Incomplete $^{15}N/^{13}C$ labeling led to residual intramolecular NOE cross-peaks, which were distinguished using control spectra of apo-NDM-1.

**Molecular Docking and Dynamics Simulations of the Thanatin–NDM-1 Complex**

**Protein preparation:*

The initial structure of NDM-1 was obtained from the Protein Data Bank (PDB ID: pdb_00005ZR8). All residues with multiple rotamers were inspected manually, and the conformations with the highest occupancy were retained. The sulfate ion present in the crystal structure was removed, while the two $Zn^{2+}$ ions located in the active site were retained. Hydrogen atoms were added using UCSF Chimera. Protonation states of residues within 2 Å of the zinc ions were carefully evaluated based on their coordination geometry and local environment. The following protonation states were assigned: Asp124 ($OD2^-$), His120 (HID), His122 (HIE), His189 (HID), His250 (HID), and Cys208 (deprotonated). The sequence used for NMR experiments differs by 25 residues from the crystallographic construct (PDB 5ZR8), primarily located at the N-termini. These regions are structurally disordered and distant from the catalytic site, and thus were not expected to affect the active-site geometry or the overall coordination environment of the $Zn^{2+}$ ions.The net charge of the zinc  site was verified to ensure a physically consistent metal–ligand coordination environment prior to system solvation.

**HADDOCK Docking:*

Molecular docking of Thanatin-NDM-1 was performed using HADDOCK 2.4 with the OPLSX force field [3,4]. The NMR structure of thanatin (PDB ID: pdb_00006aab) was used as the ligand for all simulations. NMR chemical-shift perturbations (CSPs) and intermolecular NOEs were used to define ambiguous interaction restraints (AIRs). All thanatin residues were treated as active to allow full sampling of its compact β-hairpin surface. NDM-1 active residues were defined as those with CSPs > 0.017 ppm (residues 45–48, 73, 94, 97, 122–123, 151–155), while passive residues were assigned automatically by HADDOCK as solvent-accessible neighbors within 6.5 Å. The docking protocol proceeded in three stages: (1) randomization of orientations and rigid-body energy minimization (1,000 models); (2) semi-flexible simulated annealing in torsion angle space (200 models); and (3) final refinement in explicit solvent (200 models). Histidine protonation states were explicitly defined to reflect experimental pH 7 (e.g., His120, His133, His189, His228, and His250 as Nδ-protonated; His61, His122, His159, and His261 as

Nε-protonated). Clustering was performed using an interface-RMSD cutoff of 5.0 Å and a minimum cluster size of 4. Clusters were ranked by the HADDOCK score, calculated as: $1.0*E_{vdw} + 0.2 * E_{elec} + 1.0 * E_{desolv} + 0.1 * E_{air}$. The final cluster selected for further analysis combined the lowest HADDOCK score, largest cluster size, and best agreement with experimental CSP/NOE data. Detailed docking statistics, energy terms, and cluster tables are provided in Table SI1.

**Molecular dynamics (MD) simulations:*

MD simulations were performed to investigate the structural dynamics of NDM-1 in its apo-form and in complex with the antimicrobial peptide Thanatin. All simulations were carried out using GROMACS 2024.3 [5,6] with the AMBER99SB-ILDN force field and the SPC/E explicit water model. The binuclear zinc active site was maintained using harmonic distance restraints between $Zn^{2+}$ ions and coordinating residues (His120, His122, His189, Cys208, Asp124, and His250) [7] with force constants of 10,000 $kJ \cdot mol^{-1} \cdot nm^{-2}$, together with an inter-zinc restraint of 5,000 $kJ \cdot mol^{-1} \cdot nm^{-2}$.

Each system was solvated in a periodic cubic box with a minimum solute–box edge distance of 1.0 nm, neutralized with counterions, and adjusted to 0.15 M NaCl [8]. Energy minimization was performed using the steepest descent algorithm. Systems were equilibrated in the NPT ensemble for 500 ps at 300 K using the velocity-rescaling thermostat (τ□ = 0.1 ps) applied separately to solute and solvent groups, and at 1 bar using the Parrinello–Rahman barostat (τ□ = 5.0 ps; compressibility = $4.5 \times 10^{-5}$ $bar^{-1}$). During equilibration, positional restraints (1000 $kJ \cdot mol^{-1} \cdot nm^{-2}$) were applied to heavy atoms of the protein, peptide, and zinc ions.

System density stabilized within the first 200 ps of NPT equilibration and remained constant thereafter, with pressure fluctuations centered at 1 bar without systematic drift. The final equilibrated box volume remained stable, indicating convergence of bulk solvent properties. Following stabilization of density and box dimensions, production simulations were performed in the NVT ensemble to maintain fixed-volume conditions for consistent comparison between independent trajectories.

Production simulations employed a 2 fs time step with the leap-frog integrator. All bonds involving hydrogen atoms were constrained using the LINCS algorithm. Long-range electrostatics were treated using the Particle Mesh Ewald (PME) method with a 1.0 nm real-space cutoff, and van der Waals interactions were truncated at 1.0 nm with analytical dispersion corrections applied to energy and pressure. Coordinates were saved every 2 ps.

For each state (apo and peptide-bound), one 150 ns production trajectory was generated, together with five independent 20 ns trajectories initiated from the equilibrated structure with randomized initial velocities, yielding a total sampling time of 250 ns per state. Center-of-mass motion was removed linearly. Structural analyses (RMSD, per-residue RMSF, and metal–coordiation distances [7]) were performed using standard GROMACS tools. Visualization was conducted in PyMOL. All simulations were executed on the UCR High-Performance Computing Cluster (HPCC).

**CD Thermal Denaturation**

**Protein preparation:*

NDM-1 was expressed and purified as described in the main text. *Apo preparation:* apo-protein was generated by inclusion of 10 mM EDTA during TEV cleavage (overnight, 4 °C) followed by $Ni^{2+}$ affinity purification and buffer exchange to remove residual metal and EDTA. *Zn reconstitution:* Zn-bound samples were prepared by adding $ZnCl_2$ stock to give 4.0 molar equivalents per monomer, incubating 30 min at 25 °C, and buffer-exchanging into 10 mM

HEPES, 100 mM NaCl, pH 7.5. Thanatin was purchased (NovoPro; supplied without TFA) and was prepared as a 1 mg·mL$^{-1}$ stock in Milli-Q water and added at the indicated stoichiometry. Samples were adjusted to 35 µM for CD measurements.

**CD measurement:*

Thermal denaturation experiments were performed on a Jasco J-1500 spectropolarimeter equipped with a Peltier PTC-517 temperature controller. Spectra were recorded in 1 mm quartz cuvettes at 222 nm from 20 to 85 °C with a heating rate of 1 °C·min$^{-1}$. Wait 50 s at each temperature. Acquisition parameters were: bandwidth 1 nm, response (D.I.T.) 2 s, data pitch 1 nm, scan speed 50 nm·min$^{-1}$, and 1 cycle. Buffer baselines were recorded and subtracted from sample traces.

**Data analysis:*

Raw ellipticity values were normalized to the fraction unfolded (fd) by setting pre-transition baselines to 0 and post-transition plateaus to 1. Normalized data were fitted to a Boltzmann sigmoidal function,

$$fd = 1 / (1 + exp[(Tm – T)/dT])$$

where $T_m$ is the apparent melting temperature and dT is the slope factor (transition width). Fits were performed in OriginPro, and reported values represent mean ± SD from three independent replicates. Statistical comparisons were made using two-sample t-tests; differences ≤1 °C were considered below assay resolution. All raw traces, replicate datasets, and fit parameters are provided in the Supporting Information (Figures SI2, Tables SI2).

**Bacterial Strains and Culture Conditions**

*Escherichia coli* BL21(DE3) cells harboring pET28a(+)-TEV:NDM-1 were used for expression and growth assays. Negative controls were wild-type *E. coli* BL21 and BL21 transformed with empty pET28a(+)-TEV. Overnight cultures were grown in LB medium at 37 °C with orbital shaking (250 rpm). Antibiotic selection was applied as follows: NDM-1 and empty-vector strains, kanamycin (50 µg $mL^{-1}$); TEV control (where used), ampicillin (100 µg $mL^{-1}$).

**Growth Synergy and AUC Measurements**

Overnight cultures of *E. coli* BL21(pET28a-TEV:NDM-1), wild-type BL21, and empty-vector controls were grown in LB + kanamycin (50 µg $mL^{-1}$), then subcultured 1:100 into fresh LB + IPTG (1 mM) and grown 4–5 h at 37 °C prior to normalization. Synergy assays were performed in GenClone 96-well plates (200 µL final volume/well, $n$=8 technical replicates/condition unless noted, including wild-type/vector controls). NDM-1 expression was induced with IPTG (1 mM); thanatin (NovoPro, TFA-free) was used at 6 µg $mL^{-1}$ with imipenem varied per assay requirements.

Cultures were normalized to $OD_{600}$ = 1.0 in the Synergy HTX plate reader (BioTek; Gen5 software; raw absorbance, no pathlength correction), and 2 µL/well was inoculated immediately. Plates (with lids) were incubated at 37 °C with orbital shaking (180 cycles $min^{-1}$, 6 mm) for 8 h. PBS (180 µL) was added to unused wells to prevent edge effects. $OD_{600}$ was recorded at intervals.

Area Under the Curve (AUC) was calculated in Prism v9.0.0 (trapezoidal rule, baseline = 0); outliers were excluded by ROUT (Q = 1%). AUC was analyzed by unpaired $t$-tests; terminal $OD_{600}$ by Mann-Whitney $U$. Exact $p$ values and $n$ definitions are in figure legends. Wild-type/vector controls were included throughout.

Reagents. Thanatin (NovoPro, TFA free) and imipenem; IPTG (Sigma I6758), kanamycin sulfate (Fisher BP906), and ampicillin (Fisher BP1760) were used as received. Stocks in sterile water were stored at −20 °C and used within 2 weeks. Kanamycin (50 µg $mL^{-1}$) maintained the NDM-1 plasmid.

**Viable Cell Counting (CFU/mL)**

At 8 h, 100 µL culture aliquots were serially diluted 10-fold in sterile PBS. From each dilution, 5 µL was spotted in triplicate onto LB agar plates [NaCl (5 g $L^{-1}$), tryptone (10 g $L^{-1}$), yeast extract (5 g $L^{-1}$), agar (15 g $L^{-1}$); Fisher BioReagents; autoclaved]. Plates were dried prior to spotting and incubated statically (37 °C, 16–18 h). Colonies were enumerated from spots yielding 3–30 colonies, and CFU $mL^{-1}$ values (with % expansion) were calculated relative to $T_0$ inoculum. Data were analyzed by Mann–Whitney *U* tests.

### ➢ References

(1) Vranken, W. F.; Boucher, W.; Stevens, T. J.; Fogh, R. H.; Pajon, A.; Llinas, M.; Ulrich, E. L.; Markley, J. L.; Ionides, J.; Laue, E. D. The CCPN Data Model for NMR Spectroscopy: Development of a Software Pipeline. *Proteins Struct. Funct. Bioinforma.* **2005**, *59* (4), 687–696. https://doi.org/10.1002/prot.20449.

(2) Herrmann, R.; García-García, F. J.; Reller, A. Rapid Degradation of Zinc Oxide Nanoparticles by Phosphate Ions. *Beilstein J. Nanotechnol.* **2014**, *5*, 2007–2015. https://doi.org/10.3762/bjnano.5.209.

(3) Honorato, R. V.; Koukos, P. I.; Jiménez-García, B.; Tsaregorodtsev, A.; Verlato, M.; Giachetti, A.; Rosato, A.; Bonvin, A. M. J. J. Structural Biology in the Clouds: The WeNMR-EOSC Ecosystem. *Front. Mol. Biosci.* **2021**, *8*. https://doi.org/10.3389/fmolb.2021.729513.

(4) Vargas Honorato, R. V.; Trellet, M. E.; Jiménez-García, B.; Schaarschmidt, J. J.; Giulini, M.; Reys, V.; Koukos, P. I.; Rodrigues, J. P. G. L. M.; Karaca, E.; van Zundert, G. C. P.; Roel-Touris, J.; van Noort, C. W.; Jandová, Z.; Melquiond, A. S. J.; Bonvin, A. M. J. J. The

HADDOCK2.4 Web Server for Integrative Modeling of Biomolecular Complexes. *Nat. Protoc.* **2024**, *19* (11), 3219–3241. https://doi.org/10.1038/s41596-024-01011-0.

(5) Abraham, M. J.; Murtola, T.; Schulz, R.; Páll, S.; Smith, J. C.; Hess, B.; Lindahl, E. GROMACS: High Performance Molecular Simulations through Multi-Level Parallelism from Laptops to Supercomputers. *SoftwareX* **2015**, *1–2*, 19–25. https://doi.org/10.1016/j.softx.2015.06.001.

(6) Páll, S.; Zhmurov, A.; Bauer, P.; Abraham, M.; Lundborg, M.; Gray, A.; Hess, B.; Lindahl, E. Heterogeneous Parallelization and Acceleration of Molecular Dynamics Simulations in GROMACS. *J. Chem. Phys.* **2020**, *153* (13), 134110. https://doi.org/10.1063/5.0018516.

(7) Eshtiwi, A. A.; Rathbone, D. L. A Modified Bonded Model Approach for Molecular Dynamics Simulations of New Delhi Metallo-β-Lactamase. *J. Mol. Graph. Model.* **2023**, *121*, 108431. https://doi.org/10.1016/j.jmgm.2023.108431.

(8) Chen, J.; Chen, H.; Shi, Y.; Hu, F.; Lao, X.; Gao, X.; Zheng, H.; Yao, W. Probing the Effect of the Non-Active-Site Mutation Y229W in New Delhi Metallo-β-Lactamase-1 by Site-Directed Mutagenesis, Kinetic Studies, and Molecular Dynamics Simulations. *PLOS ONE* **2013**, *8* (12), e82080. https://doi.org/10.1371/journal.pone.0082080.
